# Hippocampal consummatory reward ensembles dynamically engage theta-SWR states during spatial working memory

**DOI:** 10.64898/2026.06.01.729460

**Authors:** Yoshiyuki Omura, Jun Yamamoto, Takashi Kitamura

## Abstract

Reward consumption consists of temporally structured sensory and behavioral episodes critical for memory-guided behavior. However, how hippocampal circuits organize consummatory episodes and their associated oscillatory activity remains poorly understood. Recording dorsal hippocampal CA1 activity during a delayed nonmatch-to-place T-maze task, we identified a sparse population of reward-ensemble neurons with sustained firing aligned to distinct phases of reward consumption and spatially centered on reward locations. Reward consumption was accompanied by chewing-linked, atropine-sensitive 6-Hz theta oscillations that coexisted with intermittent sharp-wave ripples (SWRs). Notably, coupling of reward-ensemble spiking to theta and SWRs differed across sample and test phases of the task, suggesting state-dependent coordination of hippocampal reward ensembles during spatial working memory. Together, these findings identify reward consumption as a distinct hippocampal state characterized by coordinated theta-SWR dynamics during mnemonic behavior.

## Introduction

Animals must remember not only where food can be found, but also the sensory and behavioral context surrounding reward consumption to guide flexible future navigation. Such representations linking spatial location with consummatory experience are critical for survival and are thought to have exerted strong evolutionary pressure on hippocampal memory systems ^1^. Consistent with this view, the hippocampus has long been implicated in representing reward locations and guiding future foraging behavior, with classic work demonstrating place-specific activity and reward-related remapping in hippocampal circuits ^2^. More recent studies, including large-scale two-photon imaging of CA1 populations, have shown that subsets of hippocampal neurons preferentially encode reward locations within learned environments ^3^. However, effective spatial working memory requires more than a static representation of reward location; it depends on encoding ongoing sensory experience and associating that experience with behavioral outcomes across temporally structured episodes of reward intake.

In support of this role, pyramidal neurons in dorsal hippocampal CA1 respond to both spatial and non-spatial sensory inputs, as well as conjunctive information linking multiple task variables ^4–7^. Rather than encoding static features, CA1 neuronal activity is organized into firing sequences that reflect the temporal structure of ongoing experience, a hallmark of hippocampal cognitive processing ^8–10^. Feeding behavior provides a natural ethological context in which such temporal organization may be especially prominent. For species that consume food in discrete bouts, feeding consists of distinct behavioral phases, including food acquisition, consummation, and termination ^11^. Despite this clear temporal structure, prior work has largely emphasized spatial correlates of reward, leaving unresolved how hippocampal activity organizes the temporal structure of consummatory episodes themselves.

A key unresolved issue is how this temporally structured organization of consummatory episode is implemented in hippocampal circuits and coordinated with state-dependent oscillatory activity. Hippocampal activity transitions across distinct oscillatory modes depending on behavioral state; theta oscillations predominate during active exploration and memory encoding, whereas sharp-wave ripples (SWRs) emerge during immobility and associative memory processing ^12,13^. Reward consummation, however, represents a unique behavioral state in which animals are largely immobile yet continue to engage in ongoing sensory sampling and consummatory actions, and has been identified as a prominent behavioral context in which SWRs are expressed ^14^. This raises the possibility that hippocampal network coordination during reward consumption differs from that observed during exploration or rest.

From this perspective, hippocampal oscillatory activity during reward consumption provides a window into how sensory and memory-related processes are coordinated in ongoing behavior. Atropine-sensitive type 2 theta oscillations, long observed during sensory-associated immobility ^15,16^, may represent a distinct hippocampal state during consummatory behavior. However, hippocampal oscillatory states during reward consumption remain poorly characterized. Moreover, how reward-related hippocampal neuronal populations participate in these oscillatory states as behavioral and mnemonic demands change during spatial working memory remains largely unexplored.

At the circuit level, the radial organization of CA1 provides a substrate for integrating distinct oscillatory inputs during behavior ^17^. Deep and superficial pyramidal neurons differ in their sensory and spatial coding properties ^18,19^, while entorhinal, CA2, and CA3 contribute differentially to theta- and SWR-associated activity ^20–23^. Theta-associated sensory inputs and SWR-associated excitatory activity can converge in CA1 during reward-related behaviors ^24,25^. These observations suggest that reward-responsive neural ensembles in the hippocampal CA1 may engage multiple oscillatory states during spatial learning ^26,27^.

To examine how reward-related behaviors influence hippocampal activity dynamics during mnemonic processing, we recorded CA1 pyramidal spiking activity and local field potentials at reward zones during a delayed non-matching-to-place (DNMP) T-maze task, while simultaneously monitoring jaw movements associated with chewing. We identified a distinct minority of dorsal CA1 (dCA1) excitatory neurons whose activity was tightly aligned to specific consummatory acts and remained elevated throughout food intake. These neurons were robustly and selectively coupled to both atropine-sensitive theta and SWR oscillations. Notably, reward-ensemble coupling to these oscillatory states differed across the sample and test phases of the task, operationally defined trial epochs with distinct mnemonic demands in spatial working memory.

## Result

### Persistent spiking activity in dorsal CA1 during reward consumption

Neural activity was recorded from dorsal CA1 (dCA1) in five mice performing a delayed nonmatch-to-place (DNMP) T-maze task to assess spatial working memory (Fig. 1A; Supplementary Fig. 1). Recordings were obtained across daily sessions from mice implanted with tetrode microdrives targeting dCA1, and neurons (units) were pooled across sessions for subsequent analyses. One mouse tested under a different reward modality was excluded from the main cohort analyses unless otherwise noted. Mice were trained to alternate between sample (encoding) and test (retrieval) trials (Fig. 1B). To examine consummatory behavior, side-mounted cameras recorded jaw movements and three behavioral events: reward grasp, chewing onset, and forepaw contact with the floor (Fig. 1C; Supplementary Fig. 1K). Chewing onset and forepaw contact latencies were similar across left and right choices (Fig. 1D). Trials in which food was dropped during consumption were excluded unless otherwise noted.

**Figure 1.**
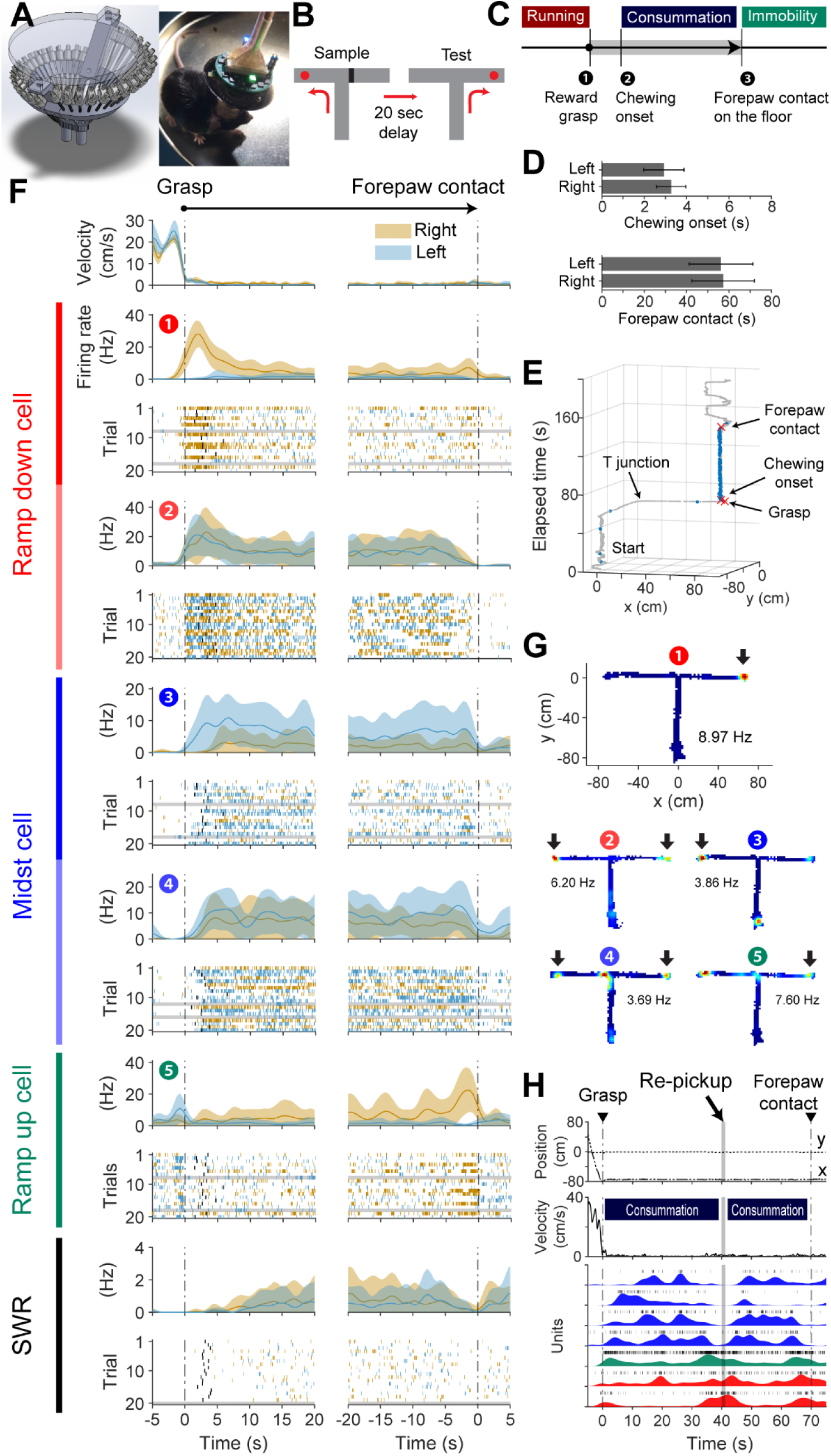
Persistent firing patterns of dorsal CA1 during reward consumption. **(A)** 34-tetrode microdrive design and implanted freely moving mouse. **(B)** DNMP T-maze task. Sample and test trials were separated by a 20 s delay. Red dots represent food reward, and the black bar is a guillotine door. **(C)** Schematic timeline of consummatory behavior. Reward intake was defined based on three behavioral timestamps: 1) reward grasp, 2) chewing onset, and 3) forepaw contact. The consummatory period was defined as the interval between reward grasp and forepaw contact. **(D)** Latencies from reward grasp to chewing onset (top) and forepaw contact (bottom) for left- and right-arm trials. **(E)** Spatiotemporal trajectory of the animal overlaid with spiking activity from a representative reward unit. Gray dots indicate the animal’s position, blue dots indicate spike occurrence, and red crosses indicate reward grasp, chewing onset, and forepaw contact. **(F)** Top: Trial-averaged head velocity aligned to reward grasp (left) and forepaw contact (right). Middle: Trial-averaged firing rates and spike rasters of representative ramp-down, midst, and ramp-up reward units. Spike rasters are colored by right-arm (orange) and left-arm (blue) trials; error trials are shaded gray. Black vertical lines indicate chewing onset. Bottom: Trial-averaged sharp-wave ripple (SWR) rates and event rasters. Solid lines indicate mean values and shaded regions indicate ±1 standard deviation. **(G)** Firing rate maps of the representative units shown in (F). Circled numbers correspond to the units shown in (F). Color indicates occupancy-normalized firing rate. Arrows denote firing fields localized to reward locations during consummation. Values indicate maximum firing rate excluding postprandial periods. **(H)** Representative activities of midst, ramp-up, and ramp-down reward units during re-picking of food fragments while animals remained at the reward site. Top: animal position (x–y). Middle: head velocity. Bottom: spike activities aligned to behavioral events.

We observed that approximately 10% of dCA1 principal units exhibited persistent spike activity during reward consumption (Fig. 1E). Within this population, some units peaked near reward grasp (ramp-down), others near forepaw contact (ramp-up), whereas a third group (midst) showed sustained activity without a clear firing peak during consummation (Fig. 1F; Supplementary Fig. 3). These units were markedly less active during immobility on error trials and during the postprandial period despite comparable head velocities (Fig. 1E; Supplementary Figs. 4D–H). Their average firing rates during consummation substantially exceeded SWR event rates (Fig. 1F). Spatially, some units exhibited firing fields confined to a single reward location, whereas others were active at both reward locations. Some units additionally exhibited place fields along the running track (Fig. 1G). Notably, brief interruptions of consummation to retrieve dropped food fragments transiently re-engaged ramping activity in ramp-down and ramp-up units, whereas midst units paused firing during these events (Fig. 1H).

### Three distinct patterns of persistent activity during reward consumption

To classify reward ensemble activity patterns, we applied principal component analysis (PCA) followed by unsupervised hierarchical clustering to binned spike counts across the consummatory period. Only excitatory units with a mean firing rate exceeding 1 Hz during consummation were included, excluding spikes occurring within SWRs to isolate persistent firing patterns independent of SWR-evoked activity. Sample and test trials were pooled because firing patterns were similar across trial types (Figs. 2F–G; Supplementary Fig. 2). To account for spatial selectivity, units were classified separately for left- and right-arm trials. This analysis revealed three distinct firing patterns, ramp-down, midst, and ramp-up, defined by their temporal firing dynamics during consummation (Fig. 2A). Similar subtype proportions were observed across left-and right-arm trials and across animals (Fig. 2B; Supplementary Fig. 2). Comparison against a shuffled null distribution revealed greater cross-arm subtype consistency than expected by chance, particularly for midst units (Figs. 2C–E; shuffle-based permutation analysis; Supplementary Table 1). Spatial selectivity, quantified as the normalized firing rate difference between choice arms during consummation, was high in ramp-down and ramp-up units but substantially lower in midst units, whereas trial selectivity between sample and test trials was weak across all subtypes (Figs. 2F–G; two-way ANOVA; Supplementary Table 1). All subtypes were distributed throughout dorsal CA1, although highly spatially selective ramp-down and ramp-up units preferentially clustered in proximal dCA1 (Fig. 2H; Supplementary Fig. 3).

**Figure 2.**
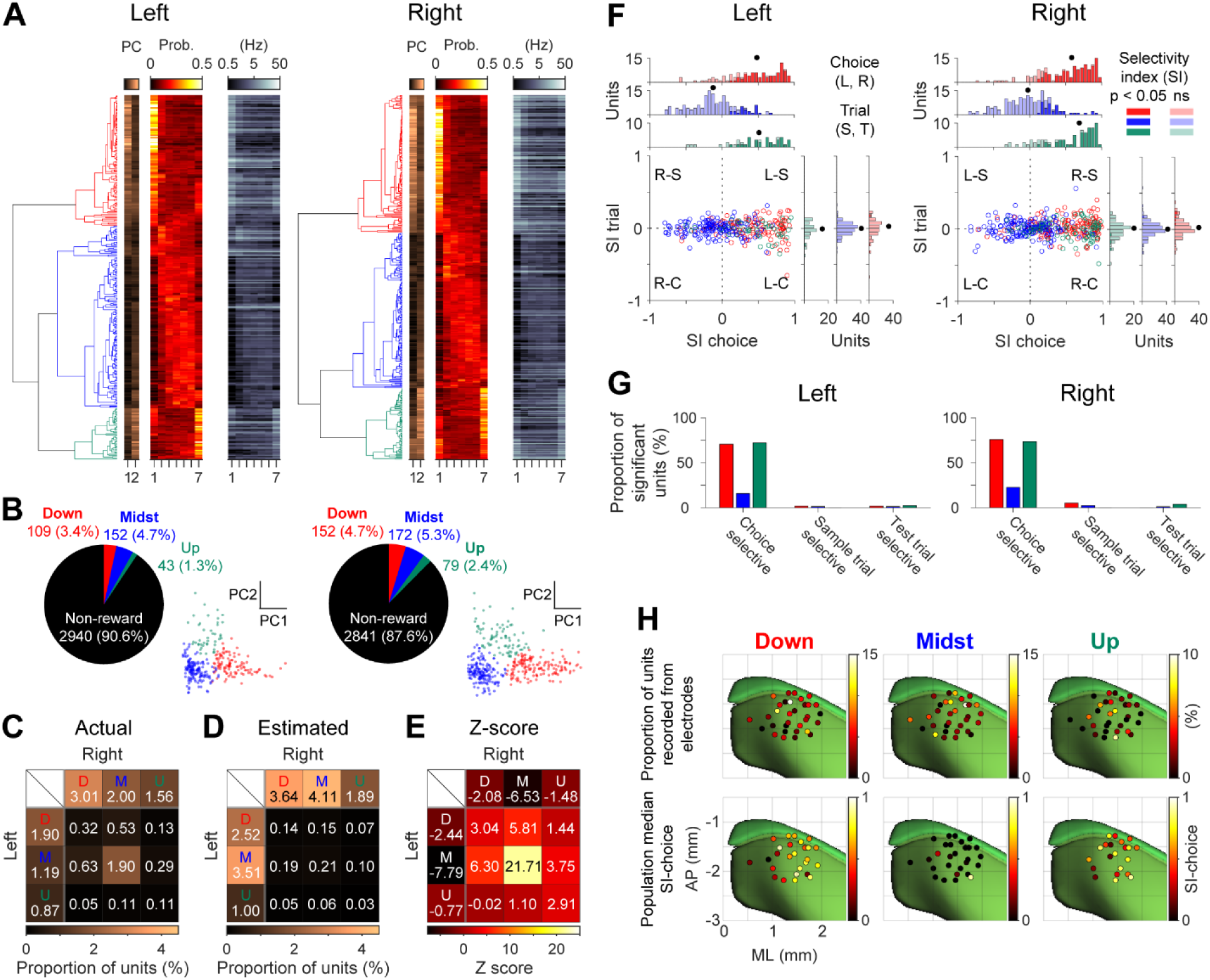
Three distinct patterns of persistent activity during reward consumption. **(A)** Hierarchical clustering of reward unit activity during consummation for left- and right-arm trials. Heatmaps show principal component (PC) scores, normalized firing rates (unit-wise normalized to sum to one), and firing rates (Hz), aligned to dendrogram order. **(B)** Proportions of ramp-down, midst, and ramp-up subtypes in left- and right-arm reward units. Right, PCA projection of clustered units. **(C)** Cross-classification of reward unit subtypes between left- and right-arm trials. Interior cells of the 4×4 matrix represent units that were reward-responsive on both arms, grouped by left-right subtype combinations. Left and upper edge boxes indicate units that were reward-responsive on only one arm. Values indicate proportions relative to all excitatory units. **(D)** Shuffled null distribution of left-right reward subtypes generated by randomly permuting subtype identities on one arm across excitatory units, assuming no correlation in subtype identity between arms. **(E)** Z-scored deviation of observed subtype distributions relative to the shuffled null distribution. **(F)** Spatial and trial selectivity of reward units. Scatter plots show spatial selectivity index versus trial selectivity index for individual units colored by subtype. Marginal histograms show subtype-specific distributions, with darker shading indicating significant selectivity. Left- and right-arm units are shown separately. **(G)** Proportion of significant spatial and trial selective units. **(H)** Anatomical distribution of reward unit subtypes in dCA1. Top panels show subtype proportions across recording locations, and bottom panels show median spatial selectivity indies across locations. Left and right recording sites are pooled with mirrored left-side locations. Statistical analyses are summarized in Supplementary Table 1.

We next examined peak firing times relative to consummatory behavioral events (Supplementary Figs. 4A–C). Ramp-down units peaked near reward grasp, whereas ramp-up units peaked near forepaw contact. In contrast, midst units increased firing after chewing onset and ceased firing before forepaw contact (Supplementary Fig. 4A). Normalized median firing time differed markedly across subtypes (Supplementary Fig. 4C; Supplementary Table 1). Together, these results demonstrate that reward ensemble subtypes exhibit distinct SWR-independent persistent firing dynamics aligned to specific phases of consummatory behavior.

To assess sensory drive, we compared average firing rates during consummation with those during immobility on error trials and during postprandial immobility (Supplementary Figs. 4D-H; Supplementary Table 1). Quantification using a preference index comparing correct and error trials revealed strong correct-biased activity in ramp-down and midst units, whereas ramp-up units showed weaker modulation. We next compared firing during consummation with postprandial immobility. Ramp-down units exhibited sustained consummation-biased firing, whereas midst units showed a brief early postprandial bias, likely reflecting residual oral processing, before shifting toward consummation-related firing. In contrast, ramp-up units exhibited stronger postprandial activity that gradually decayed over time. Together, these results indicate that reward ensemble subtypes differentially track the temporal progression of consummatory behavior.

### Reward ensembles are strongly activated by SWRs

As SWRs frequently occurred during reward consumption ^14^, we next examined reward ensemble activity during these events. Although all reward ensemble subtypes exhibited robust persistent firing outside SWRs, their activity during SWRs was predominantly excitatory, with most units strongly activated (SWR-positive; P-units) and a smaller fraction transiently inhibited (SWR-negative; N-units) (Figs. 3A-E; Supplementary Fig. 5). Peak SWR-evoked firing rates in reward ensembles substantially exceeded those of non-reward excitatory units, despite only ∼10% of total spikes occurring within SWRs (Figs. 3D and 3F). Thus, SWRs represent a temporally sparse but high-gain modulatory mode for reward ensembles. The proportions of P- and N-units varied across subtypes, with ramp-down and ramp-up units more frequently classified as P-units than midst units, irrespective of spatial selectivity (Fig. 3E; Supplementary Fig. 5).

**Figure 3.**
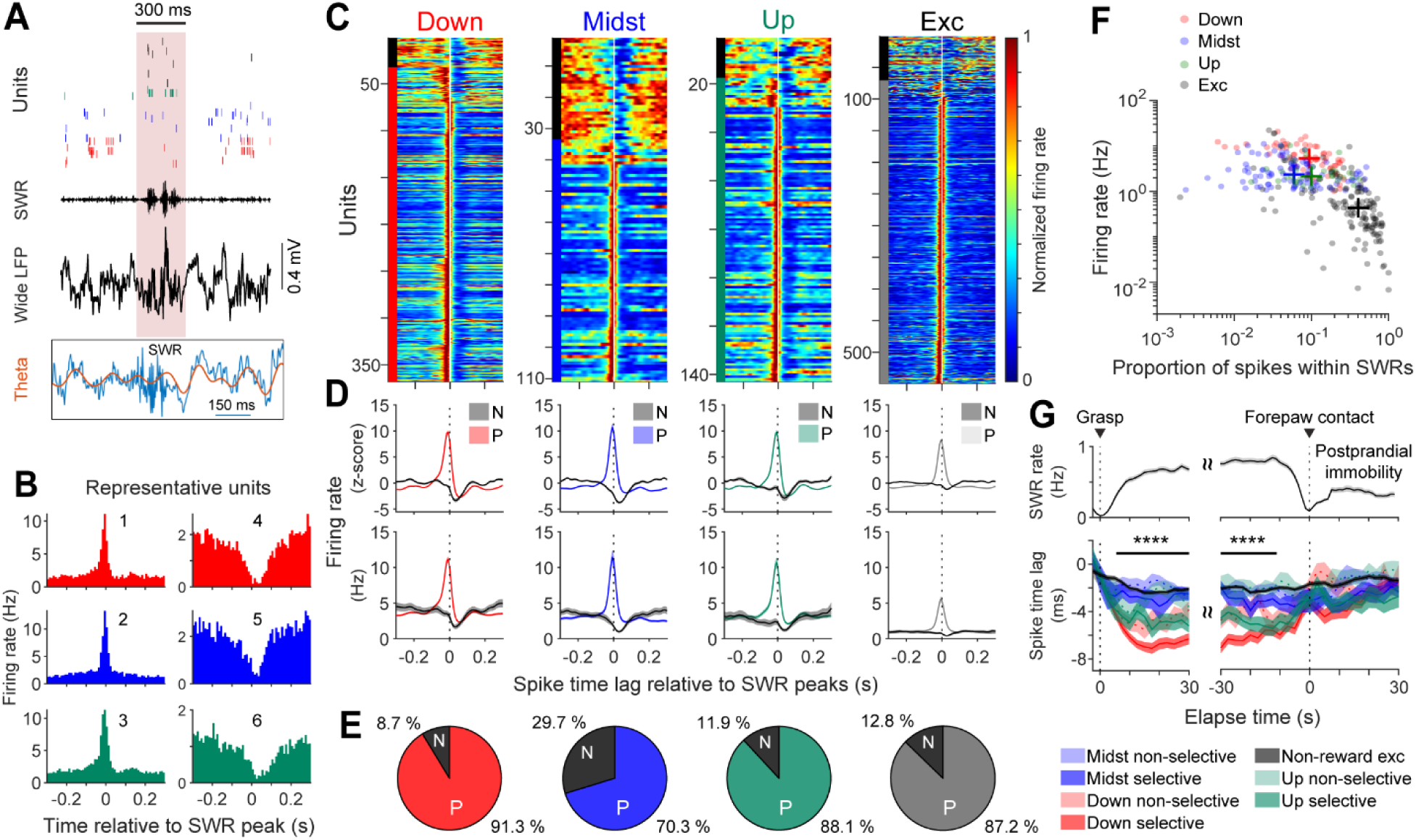
Reward ensemble activity during SWRs in consummation. **(A)** Reward ensemble activity aligned to SWR events. Representative spike rasters from reward and non-reward excitatory units are shown together with wideband, SWR-band-pass-filtered, and theta-band-pass-filtered LFP traces aligned to the SWR peaks. Shading denotes the SWR window. Red, ramp-down; blue, midst; green, ramp-up; black, non-reward excitatory units. **(B)** Representative peri-SWR firing histograms from SWR-positive (P-units: 1–3) and SWR-negative (N-units: 4–6) reward units during reward consumption. **(C)** Peri-SWR firing histograms of spatially selective reward and non-reward units aligned to SWR peaks (white vertical line). Firing rates were normalized to the maximum firing rate of each unit. Color bars on the left indicate P-units and N-units (black). **(D)** Population averaged z-scored peri-SWR firing histograms (top) and mean firing rates (bottom). Shaded regions indicate ± s.e.m. **(E)** Proportion of P- and N-units within each spatially selective reward subtypes and non-reward excitatory units. **(F)** Relationship between averaged firing rate and proportion of spikes within SWRs during reward consumption. Each dot represents a single unit, with color denoting reward subtypes and non-reward units. **(G)** Time-resolved population averaged spike time lag relative to SWR peaks. Horizontal black lines denote time points at which one-way ANOVA across reward subtypes and non-reward excitatory units reached p < 0.0001, computed independently at each time point. See Supplementary Table 1 for statistical results.

We next examined SWR-triggered spike timing relative to SWR peaks (Fig. 3G). During reward consumption, spike time lags differed significantly across reward subtypes, with ramp-down and ramp-up units firing earlier relative to SWR peaks than midst and non-reward excitatory units. In contrast, subtype differences were substantially weaker during postprandial immobility. Time-resolved analyses using one-way ANOVA at each time point revealed earlier SWR-relative firing in ramp-down and ramp-up units during reward consumption, particularly among spatially selective units (Fig. 3G; Supplementary Table 1). Together, these results indicate subtype- and spatial-selectivity-dependent temporal organization of reward ensemble activity during SWRs throughout reward consumption.

### Shifting theta phase preference during REM sleep

As previous findings implicate functional differences between CA1 sublayers ^25^, we examined theta phase preference of reward ensembles across behavioral states. During REM sleep, reward units across subtypes exhibited robust theta-rhythmic spiking, with some units preferentially firing near the theta peak and others near the trough (Supplementary Fig. 6). Only units with significant phase locking in both running and REM sleep states (Rayleigh test, p < 0.05) were included in the analysis. During open-field exploration, the majority of units fired preferentially on the descending phase or near the trough of theta (Supplementary Fig. 6B). In contrast, during REM sleep, phase preferences were more broadly distributed. Although we did not observe the pronounced bimodal distribution of preferred phases during REM as reported previously ^25^, a comparison of phase preference proportions between REM sleep and running revealed a state-dependent shift. Specifically, the ratio of units preferring each theta phase (REM vs. running) peaked near the theta peak, indicating an increased recruitment of peak-preferring units during REM sleep, particularly among spatially selective subtypes (Supplementary Fig. 6B). Together with the SWR analyses, these results suggest state-dependent temporal coordination of spatially selective reward ensembles.

### Phase coherence between atropine sensitive theta activity and chewing rhythmicity

We next examined oscillatory states outside SWRs during reward consumption. As animals transitioned from running to consummatory behavior, hippocampal theta oscillations shifted from approximately 8 Hz to 6 Hz (Figs. 4A–B). Wavelet spectrograms and time-resolved spectral analyses revealed sustained rhythmic activity in the ∼6-Hz range, particularly during early consummation, together with enhanced gamma-band power (Figs. 4A–B; Supplementary Figs. 7C–E). Coherence analyses further revealed synchronized ∼6-Hz activity across recording electrodes (Supplementary Fig. 7B). In parallel, the theta-delta power ratio peaked near chewing onset and gradually declined toward forepaw contact, while SWR rates increased inversely; notably, SWRs occurred during periods of ongoing theta activity but intermittently disrupted theta oscillations, indicating a smooth yet dynamically structured transition in hippocampal network state across the consummatory epoch (Figs. 3A and 4C; Supplementary Figs. 7D–E). To assess whether this oscillatory state was associated with orofacial sensorimotor rhythmicity, we simultaneously tracked jaw movements by measuring the distance between the maxilla and mandible (Supplementary Figs. 8A–B). Jaw movement rhythmicity exhibited a spectral peak near 6 Hz, closely resembling that of hippocampal theta oscillations, and phase-phase analyses demonstrated coherent coupling between the two rhythms (Fig. 4D; Supplementary Fig. 8C).

**Figure 4.**
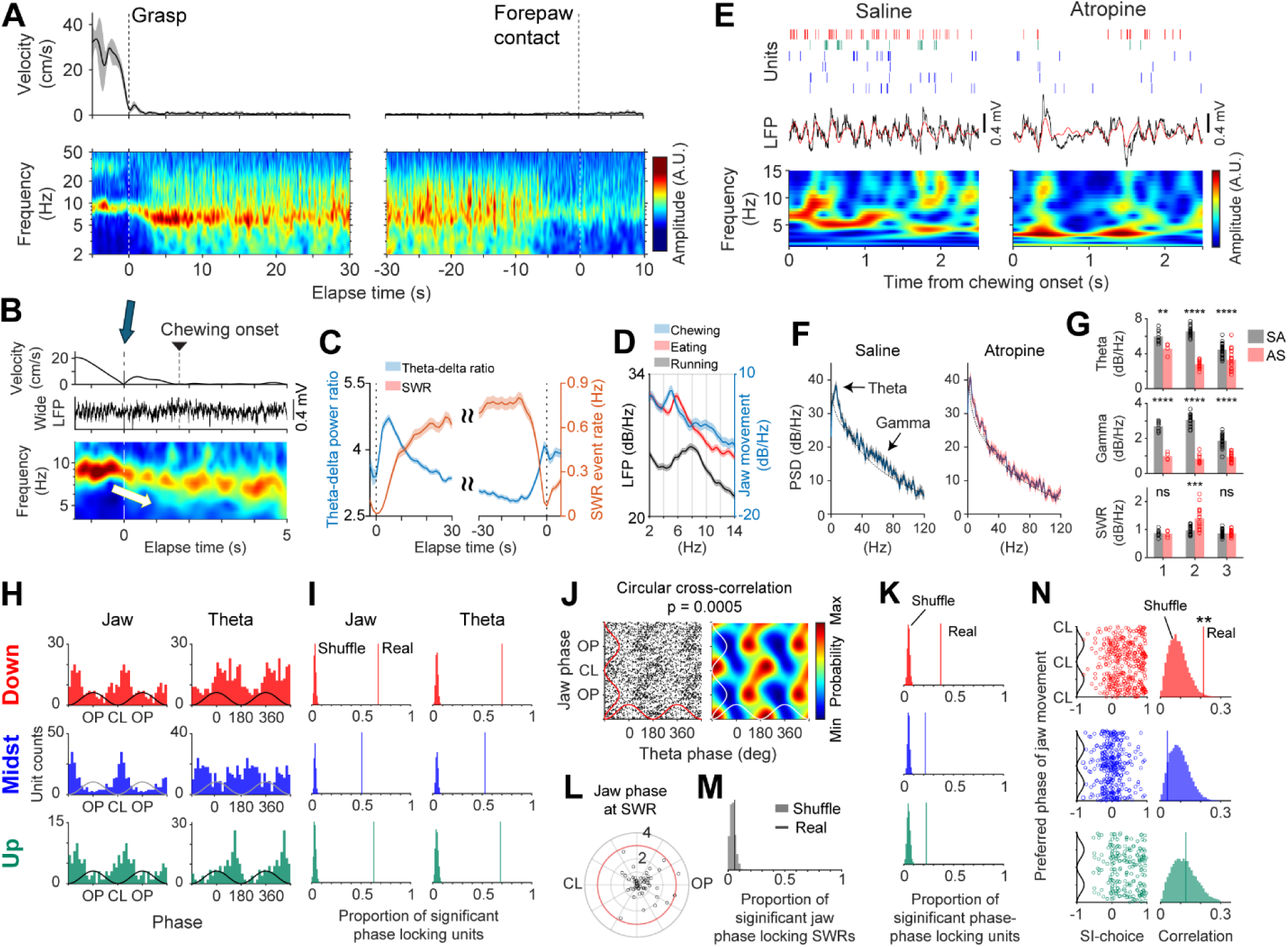
Phase synchronization of atropine-sensitive theta oscillations with chewing rhythms. **(A)** Trial-averaged head velocity and wavelet-transformed hippocampal LFP spectrograms aligned to reward grasp (left) and forepaw contact (right). **(B)** Magnified view of LFP transition aligned to reward grasp, highlighting the shift in theta frequency corresponding to the behavioral transition from running to consummation. **(C)** Theta-delta power ratio and SWR rate. **(D)** LFP power spectral density (PSD) during running and consummatory states (left axis), together with PSD of jaw movement rhythmicity during chewing (right axis). **(E)** Representative reward-units firing activity and LFP under saline and atropine conditions. Note the lack of periodicity and firing reduction in the atropine condition. **(F)** PSD of LFP during consummation, under saline (left) or atropine (right) conditions. The 95% confidence interval is shaded. **(G)** Comparison of FOOOF-derived periodic power under saline (NS) and atropine sulfate (AS) conditions for three mice. Multiple trials were pooled within each mouse. From top to bottom: theta, gamma, and ripple-band frequency ranges. **p < 1e−2; ****p < 1e−4; ns, not significant. **(H)** Preferred phase distributions relative to jaw movement rhythmicity (left) and theta oscillations (right) for each subtype. Only significantly phase-locked units were included. **(I)** Proportion of significantly phase-locked units relative to jaw movement rhythmicity and theta oscillations compared with 2,000 spike-shuffled controls. All comparisons were significant (p < 5e-4). **(J)** Circular-circular correlation analysis of spike phases between jaw movement rhythmicity and theta oscillations. Left: example of a ramp-down unit exhibiting significant co-modulation. Right: density heatmap of spike-phase relationships between the two rhythms. **(K)** Proportion of significantly co-modulated units. Significant co-modulation was defined by circular-circular correlation between spike phases relative to jaw movement rhythmicity and theta oscillations (p < 0.05) and compared with 2,000 spike-shuffled controls. **(L)** Distribution of preferred phases of jaw movement rhythm at SWR peaks. Red circle indicates the Rayleigh significance threshold. Each dot represents one session. **(M)** Proportion of significant phase-locking between SWR peaks and jaw movement rhythmicity compared with 2,000 shuffled controls. **(N)** Relationship between jaw-rhythm phase preference and spatial selectivity across reward-unit subtypes. Each dot represents one unit. Circular-linear correlations were compared with 2,000 spike-shuffled controls. * p < 0.05, **p < 0.01. See Supplementary Table 1 for statistical results.

These observations raised the possibility that the coordinated ∼6-Hz oscillatory state observed during consummation reflects cholinergic mechanisms characteristic of type 2 theta. To test this, we administered atropine sulfate (50 mg/kg) or 0.9% saline intraperitoneally and examined the effect of muscarinic blockade on hippocampal oscillatory dynamics during consummation. Atropine markedly reduced ∼6-Hz rhythmicity and gamma power during consummation, while rhythmic activity during running remained largely intact (Figs. 4E–G; Supplementary Table 1; Supplementary Figs. 7F–I). In contrast, ripple-band power was not consistently reduced by atropine. Reward-unit firing during consummation was also reduced following atropine administration (Fig. 4E). Because chewing behavior persisted under atropine, these findings suggest that the ∼6-Hz oscillatory state observed during consummation is consistent with atropine-sensitive type 2 theta rather than chewing-related movement artifacts.

### Chewing- and theta-locked activity in reward ensembles with spatial selectivity

We next examined whether reward-unit spiking is jointly modulated by hippocampal theta phase and jaw movement rhythmicity during consummation, excluding spikes occurring within SWRs. Across reward subtypes, units were preferentially phase-locked to the mouth-closing phase of the chewing cycle (Fig. 4H). With respect to theta oscillations, ramp-down and ramp-up units were preferentially phase-locked to the descending phase, whereas midst units were preferentially locked near the theta trough (Fig. 4H). A substantial fraction of units exhibited significant phase locking to both jaw movement rhythmicity and theta oscillations compared with spike-time shuffled controls (Fig. 4I; Supplementary Table 1). Although many units were significantly phase-locked to both rhythms, no consistent phase lag between jaw movement and theta oscillations was observed at the population level (Supplementary Fig. 8D; Supplementary Table 1). We therefore quantified spike co-modulation jointly synchronized to both rhythms (Fig. 4J). Across all reward subtypes, the proportion of significantly co-modulated units exceeded shuffled controls (Fig. 4K; Supplementary Table 1). In contrast, SWR peak timing was not phase-locked to jaw movement rhythmicity, suggesting that SWR timing is largely independent of orofacial sensorimotor rhythms (Figs. 4L–M; Supplementary Table 1).

We next examined whether spatial selectivity was related to spike timing relative to jaw movement rhythmicity and theta oscillations. Correlation analyses revealed positive relationships between spatial selectivity and phase-locking strength in ramp-down and ramp-up units, whereas midst units showed little or no relationship (Supplementary Fig. 8E; circular-linear correlation analyses). Consistent with these findings, circular-linear correlation analyses further revealed subtype-specific associations between spatial selectivity and jaw-phase spike timing in ramp-down units but not in the other subtypes (Fig. 4N; Supplementary Table 1). Together, these results suggest that theta-phase organization of reward-unit spiking is associated with both consummatory sensorimotor rhythmicity and spatial selectivity in the hippocampal network.

### Persistent theta-locked firing in reward ensembles with spatial selectivity

We next asked whether theta-phase coordination of reward-unit spiking is transient or persistent throughout consummation. To address this, we quantified time-resolved spike-LFP phase coupling using pairwise phase consistency (PPC2) ^28^ and spike-phase autocorrelograms ^22^, excluding spikes occurring within SWRs.

Population-averaged PPC2 spectrograms revealed robust theta-phase locking to ∼6-Hz theta oscillations across reward subtypes (Figs. 5A–C). Theta locking persisted for tens of seconds after reward grasp, weakened during mid-consummation, and re-emerged before forepaw contact, with no sustained locking in other frequency bands (Figs. 5A–C). Spike-phase autocorrelograms further confirmed persistent theta-phase locking rather than transient phase precession (Figs. 5D–F). At reward grasp, preferred locking frequency shifted from ∼8 Hz toward ∼6 Hz across subtypes (Fig. 5H; Supplementary Table 1). Ramp-down units began firing at a late theta phase and subsequently became stably locked to the descending phase following chewing onset, whereas ramp-up units progressively advanced toward earlier theta phases near consummation termination (Figs. 5G and 5J). Throughout consummation, spatially selective units exhibited stronger theta-phase locking than non-selective units (Fig. 5I; Supplementary Fig. 9; Supplementary Table 1).

**Figure 5.**
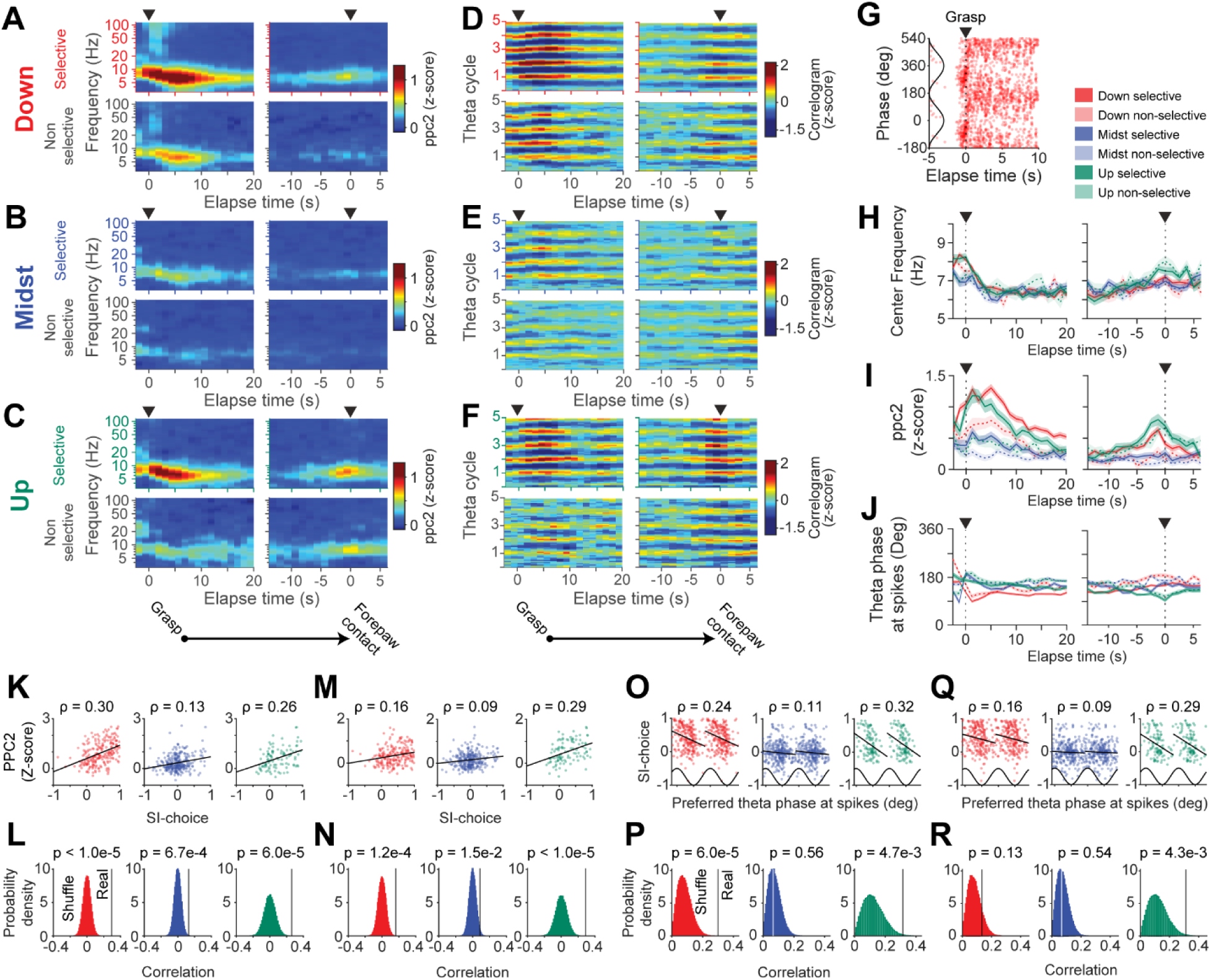
Persistent theta phase-locking correlates with spatial selectivity. **(A–C)** Population-averaged PPC2 spectrograms for ramp-down (A), midst (B), and ramp-up (C) reward subtypes. Warmer colors indicate stronger phase-locking at each frequency and time point. Top, spatially selective units; bottom, non-selective units. Left, aligned at reward grasp; right, aligned at forepaw contact. **(D–F)** Population-averaged spike phase-autocorrelograms for ramp-down (D), midst (E), and ramp-up (F) units. Warmer colors indicate higher spike probability. Layout as in A–C. **(G)** Spike-phase raster of a representative spatially selective ramp-down unit aligned to reward grasp. Spikes are plotted as a function of time and theta phase over two cycles. **(H)** Theta frequency at maximum phase-locking. Lines represent peaks of population-averaged PPC2 spectrograms, with shaded area indicating ±1 s.e.m. Left, aligned at reward grasp; right, aligned at forepaw contact. **(I)** Population-averaged PPC2 within the theta frequency range (4–10 Hz). Shaded regions indicate ±1 s.e.m. **(J)** Population-averaged preferred theta phase with ±1 s.e.m. **(K)** Relationship between spatial selectivity and theta phase-locking strength (PPC2) during the 5 s following chewing onset. Each dot represents one unit. **(L)** Comparison of correlations in K with shuffled null distributions generated by randomizing the relationship between spatial selectivity and PPC2 across units. **(M,N)** Same as K and L, respectively, for the 5 s preceding forepaw contact. **(O)** Relationship between spatial selectivity and preferred theta phase during the 5 s following chewing onset. Each dot represents one unit. **(P)** Comparison of circular-linear correlations in O with shuffled null distributions generated by randomizing the relationship between preferred theta phase and spatial selectivity across units. **(Q,R)** Same as O and P, respectively, for the 5 s preceding forepaw contact. See Supplementary Table 1 for statistical results.

Given that theta locking peaked around chewing onset and forepaw contact, we next examined the relationship between spatial selectivity and theta phase-locking during these epochs. During the 5 s following chewing onset, spatial selectivity positively correlated with theta phase-locking strength across reward subtypes, particularly in ramp-down and ramp-up units, and these relationships remained significant in permutation tests (Figs. 5K–L; Kendall’s rank correlation with permutation testing; Supplementary Table 1). Spatial selectivity was also associated with preferred theta phase in ramp-down and ramp-up units, with highly selective units preferentially firing near the theta peak (Figs. 5O–P). During the 5 s preceding forepaw contact, spatial selectivity again positively correlated with phase-locking strength, particularly in ramp-down and ramp-up units (Figs. 5M–N). Preferred theta phase was also associated with spatial selectivity during this terminal phase, most prominently in ramp-up units, with highly selective units preferentially firing near the theta peak (Figs. 5Q–R; Supplementary Table 1).

Together with the orofacial sensorimotor neuronal modulation (Fig. 4; Supplementary Fig. 8), these findings indicate that spatially selective ramping units are preferentially engaged in sustained, theta phase-locked activity across the temporal progression of reward consumption.

### Theta-gamma coupling of spikes is enhanced during sample trials

Given that spatially selective reward ensembles exhibited persistent theta-phase locking during consummatory behavior, we next examined whether these neurons also engage theta-gamma phase-amplitude coupling (TG-PAC) during spatial working memory. In the DNMP task, sample trials require encoding of the current reward location to guide subsequent reward-directed choice, thereby imposing distinct mnemonic demands relative to test trials.

During early reward consumption, LFP-based analyses revealed pronounced coupling between ∼6-Hz theta oscillations and both low- and high-gamma amplitudes (Supplementary Fig. 10E), consistent with concurrent enhancement of theta and gamma oscillatory components during consummation (Supplementary Figs. 7D–E). Spike-triggered TG-PAC analyses further revealed enhanced coupling between theta-phase and gamma-band activity across reward subtypes during the initial phase of consumption (Figs. 6A, D, and G). Because the number of units exhibiting positive TG-PAC modulation was limited, both P- and N-units were pooled for the primary analyses shown in Figure 6. PN-resolved analyses and additional temporal characterizations revealed similar overall patterns across polarity classes and gamma bands (Supplementary Figs. 10–12).

**Figure 6.**
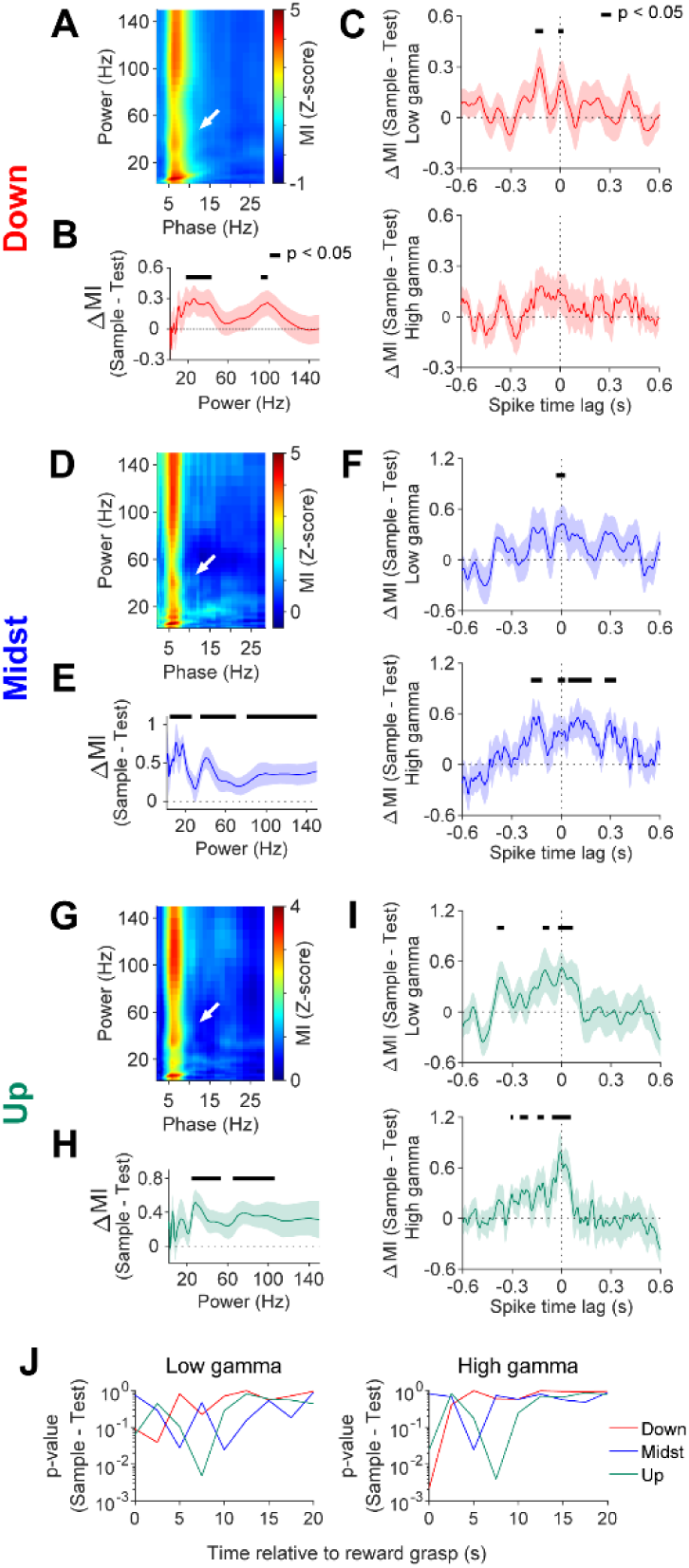
Elevated TG-PAC spike modulation during sample trials. **(A)** Population-averaged cross-frequency phase-amplitude coupling (PAC) in spatially selective ramp-down units during the 0–5 s period following reward grasp. Arrow indicates enhanced low-gamma coupling. **(B)** Trial difference in population-averaged PAC along frequency bands coupled to theta phase (4–10 Hz) in spatially selective ramp-down units. Shaded areas indicate ±1 s.e.m. Black horizontal lines indicate significant frequency ranges identified by cluster-based paired permutation testing (p < 0.05). **(C)** Trial difference in peri-spike TG-PAC between theta phase and low-gamma (top) or high-gamma (bottom) amplitude during the 0–5 s period following reward grasp. Black horizontal lines indicate significant peri-spike intervals identified by cluster-based paired permutation testing (p < 0.05). **(D–F)** Same as A–C, respectively, for spatially selective midst units during the 2.5–7.5 s period following reward grasp. **(G–I)** Same as A–C, respectively, for spatially selective ramp-up units during the 5–10 s period following reward grasp. **(J)** Time-resolved statistical significance of sample-test TG-PAC differences for low-gamma (left) and high-gamma (right) coupling across reward subtypes. Significance at each time point was assessed using paired t-tests across individual units. See Supplementary Table 1 for statistical results.

We next compared spike-based TG-PAC between sample and test trials. Spatially selective reward subtypes exhibited significantly enhanced low- and high-gamma TG-PAC during sample trials relative to test trials (Figs. 6B, E, and H; Supplementary Table 1). Peri-spike analyses further revealed that this enhancement was centered around spike timing and confined to a narrow temporal window surrounding reward grasp and chewing onset (Figs. 6C, F, I, and J; Supplementary Figs. 10–13; Supplementary Table 1). In contrast, similar trial-dependent enhancement was not observed near forepaw contact (Supplementary Fig. 12).

Together, these findings suggest that theta-gamma coupling in spatially selective reward ensembles is selectively enhanced during the early phase of reward consumption under encoding-related mnemonic demands.

### SWR-triggered spike activity is elevated during test trials

We next asked whether SWR-associated recruitment of reward ensembles differs between sample and test phases of the DNMP task. Whereas sample trials primarily require encoding of the current reward location, test trials require mapping remembered information onto the appropriate reward-directed choice, thereby imposing distinct associative demands.

Across both task phases, reward P-units exhibited robust activation around SWR peaks, substantially greater than that of non-reward excitatory units, whereas N-units showed strong suppression during SWRs irrespective of spatial selectivity (Supplementary Figs. 14A–B). Individual units maintained consistent SWR response polarity across trials, indicating stable subtype-specific recruitment during SWRs (Supplementary Fig. 16).

Having established this baseline, we next examined trial-dependent modulation of SWR-triggered firing. Spatially selective ramping P-units exhibited significantly enhanced SWR-triggered spiking during test relative to sample trials (Fig. 7A; cluster-based paired permutation tests; see also Supplementary Figs. 14A and D; Supplementary Table 1). Notably, spikes in these units occurred earlier relative to SWR peaks during test trials, with differences emerging ∼50 ms before SWR peaks (Supplementary Fig. 14C). In contrast, spatially selective midst N-units showed stronger suppression during test trials, producing greater separation between P- and N-unit activity during the test phase (Supplementary Figs. 14B and E; Supplementary Table 1). Trial-dependent modulation was substantially weaker in spatially non-selective subtypes and non-reward excitatory units.

**Figure 7.**
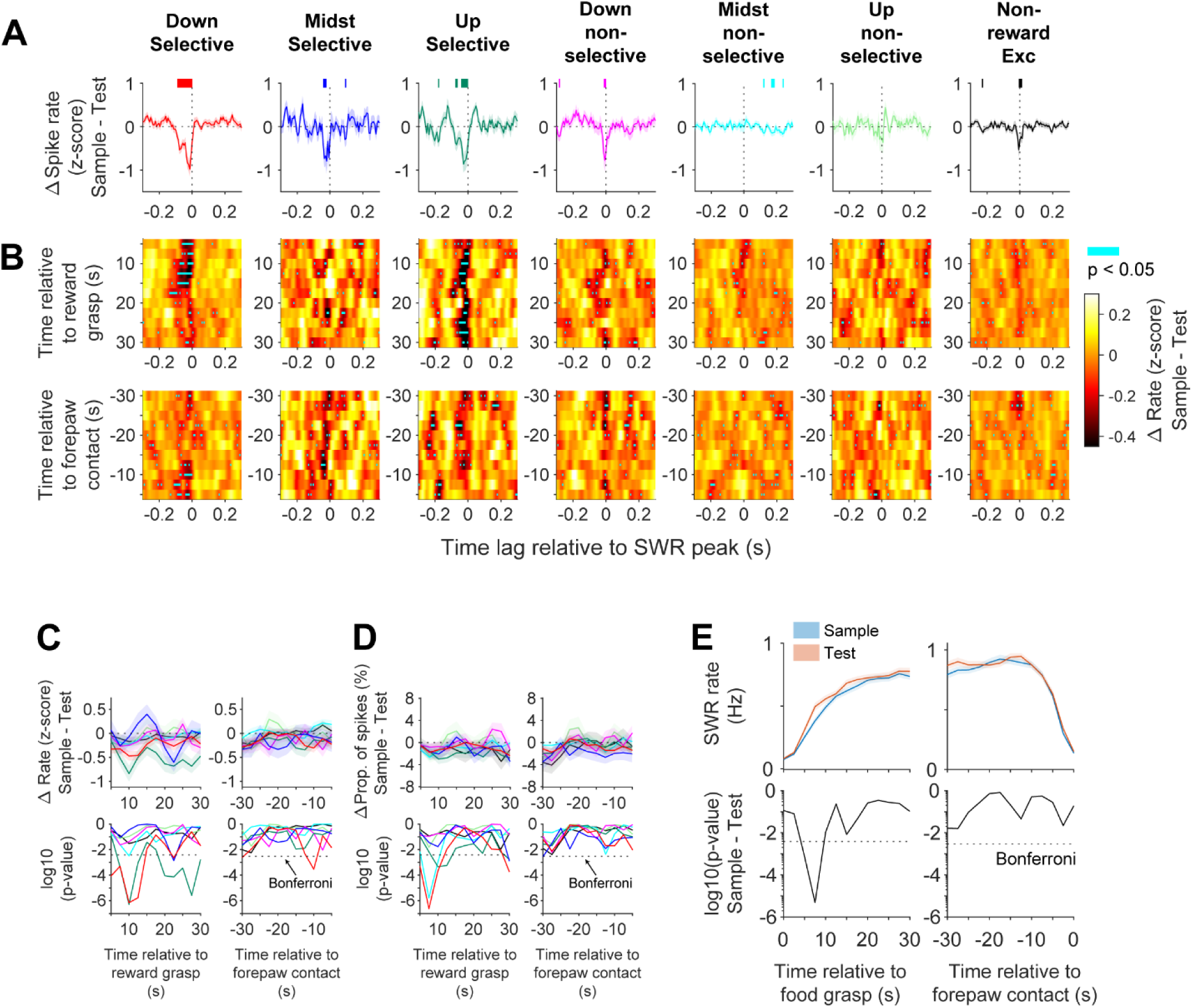
Elevated SWR-triggered spike modulation in spatially selective reward units during test trials at reward sites ipsilateral to each unit’s theta field. **(A)** Differential peri-SWR peak spike rates between sample and test trials for spatially selective and non-selective reward subtypes of P-units and non-reward excitatory P-units. Population-averaged traces are shown with shaded areas representing ±1 s.e.m. Significant differences between sample and test trials were evaluated using a cluster-based paired permutation test (500 shuffles, one-sided, p < 0.05). Horizontal lines indicate significant peri-SWR intervals. **(B)** Time-resolved differences in peri-SWR peak spike rates between sample and test trials (sample – test). Heatmaps show peri-SWR spike rate differences aligned to reward grasp (top) or forepaw contact (bottom). Cyan ticks indicate time points with significant differences determined by cluster-based paired permutation tests (500 shuffles; one-sided, p < 0.05). Darker colors indicate stronger SWR-triggered spike rates during test trials compared to sample trials. **(C)** Differences in spike rates between sample and test trials around consummatory events. Top, population-averaged spike rate differences aligned to reward grasp (left) or forepaw contact (right), plotted separately for each reward subtype. Bottom, corresponding log10(p-values) from one-sided paired t-tests at each time point. Negative Δ values indicate greater spike activity in test trials relative to sample trials. Dashed lines indicate the Bonferroni-corrected significance threshold. **(D)** Differences in the proportion of spikes occurring within SWRs between sample and test trials. Top, population-averaged changes aligned to reward grasp (left) or forepaw contact (right). Bottom, corresponding log10(p-values) from one-sided paired t-tests at each time point. Dashed lines indicate the Bonferroni-corrected significance threshold. **(E)** Comparison of SWR rates between sample and test trials. Top, Averaged SWR rates with shaded areas ±1 s.e.m. Bottom, corresponding log10(p-values) from paired t-tests at each time point. Dashed lines indicate the Bonferroni-corrected significance threshold. See Supplementary Table 1 for statistical results.

Time-resolved analyses further revealed sustained test-biased SWR-triggered spiking in spatially selective ramping P-units during the early phase of consumption, which gradually diminished later in the consummatory period (Figs. 7B–D). Although SWR event rates transiently increased during test trials shortly after chewing onset, enhanced SWR-triggered spiking persisted beyond this interval, indicating prolonged trial-dependent excitability rather than differences in SWR incidence alone (Fig. 7E). Together, these results show that recruitment of spatially selective reward ensembles during SWRs is preferentially enhanced early in reward consumption, coinciding with periods of prominent sensory-evoked 6-Hz theta oscillations (Fig. 5).

### Dual theta–SWR spiking accompanies trial-dependent SWR modulation

We next asked whether enhanced SWR-triggered firing during test trials depends specifically on reward ensembles exhibiting sustained theta-modulated spiking during consummation. To address this, we examined SWR-triggered activity at contralateral reward sites, defined as reward locations opposite to the spatial firing fields of spatially selective units. At these contralateral sites, spatially selective reward units retained SWR-locked spiking but exhibit substantially reduced 6-Hz theta modulation. As at ipsilateral reward sites, reward units at contralateral locations exhibited robust activation (P-units) or suppression (N-units) during SWRs relative to non-reward excitatory units (Supplementary Figs. 15A–B). Peak SWR-evoked firing rates during sample and test trials also remained strongly correlated at the single-unit level, indicating stable baseline SWR engagement across reward sites (Supplementary Fig. 16). In contrast to ipsilateral sites, however, trial-dependent enhancement of SWR-triggered spiking was no longer evident at contralateral locations in either P- or N-units (Fig. 8A; cluster-based paired permutation tests; see also Supplementary Figs. 15A–E; Supplementary Table 1). Spike timing relative to SWR peaks was also comparable across reward subtypes and non-reward excitatory units (Supplementary Fig. 15C). Time-resolved analyses further revealed similar SWR-triggered activity between sample and test trials throughout the consummatory period at contralateral reward sites (Figs. 8B–E; Supplementary Table 1). Therefore, these results indicate that enhanced SWR-triggered recruitment during test trials is not explained by SWR responsiveness alone, but is preferentially associated with spatially specific theta-modulated reward ensemble activity expressed at ipsilateral reward locations.

**Figure 8.**
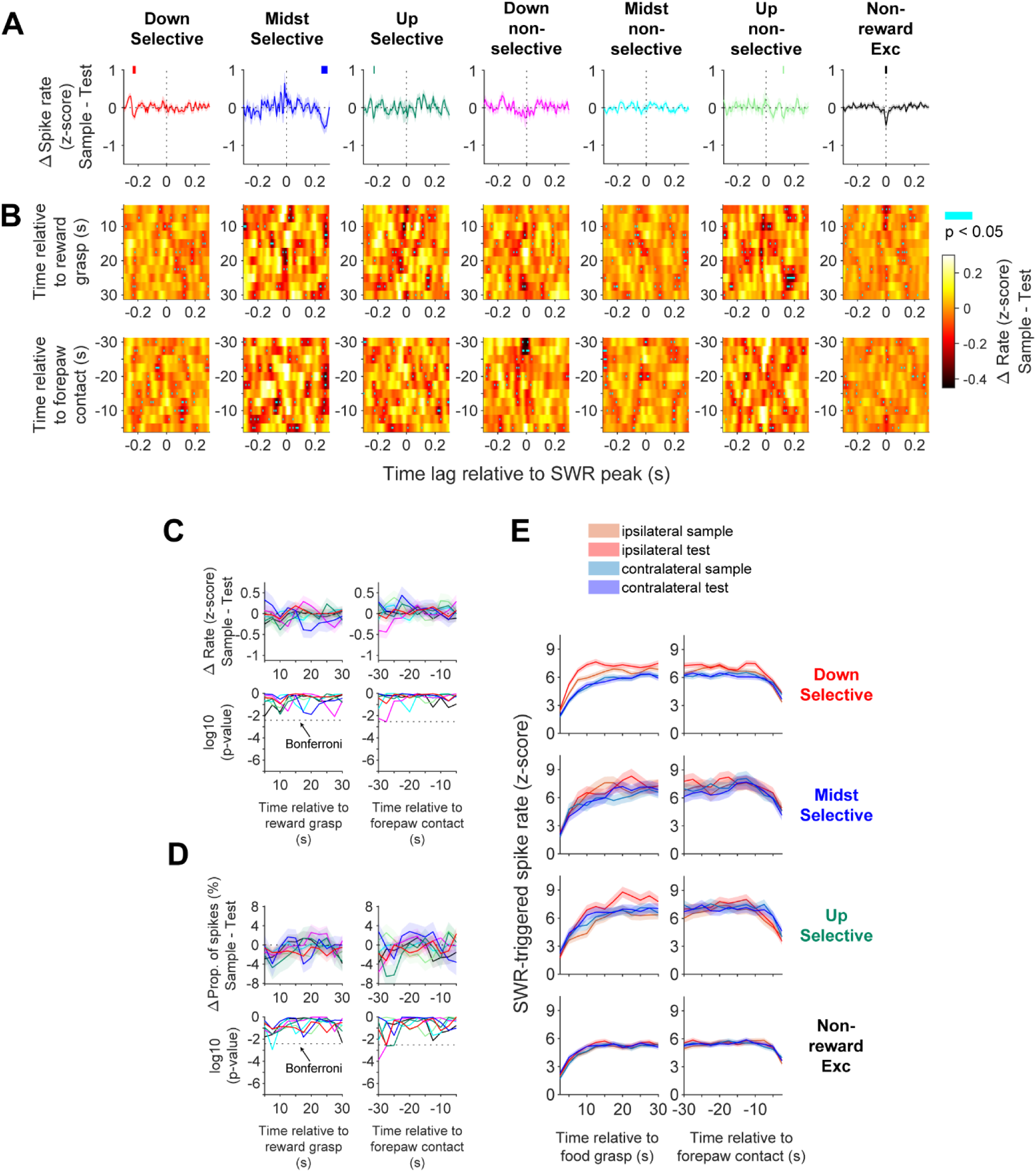
Lack of trial-dependent SWR-triggered spike modulation in spatially selective reward units at reward sites contralateral to each unit’s theta field. **(A)** Peri-SWR peak spike rates for the same spatially selective and non-selective reward P-unit subtypes and non-reward excitatory P-units shown in Figure 7A, quantified at the contralateral reward sites. Population-averaged traces are shown with shaded areas representing ±1 s.e.m. In contrast to ipsilateral reward sites (Fig. 7A), no significant trial-dependent differences between sample and test trials were observed. Significant differences were evaluated using cluster-based paired permutation tests (500 shuffles, one-sided, p < 0.05). **(B)** Heatmaps showing peri-SWR peak spike rate differences (sample – test) for the same units as in Figure 7B, quantified at contralateral reward sites. Cyan ticks denote time points with significant differences (cluster-based paired permutation tests; 500 shuffles, one-sided, p < 0.05). **(C)** Differences in spike rates between sample and test trials around consummatory events for the same reward P-unit subtypes shown in Figure 7C. Top, population-averaged spike rate differences aligned to reward grasp (left) or forepaw contact (right). Bottom, corresponding log10(p-values) from one-sided statistical comparisons at each time point. No robust trial-dependent modulation was observed across the consummatory period. Dashed lines indicate the Bonferroni-corrected significance threshold. **(D)** Differences in the proportion of spikes occurring within SWRs between sample and test trials. Top, population-averaged differences aligned to reward grasp (left) or forepaw contact (right). Bottom, corresponding log10(p-values) from one-sided statistical comparisons at each time point. Dashed lines indicate the Bonferroni-corrected significance threshold. **(E)** Time-resolved population-averaged SWR-triggered spike rates across spatially selective reward subtypes for ipsilateral sample, ipsilateral test, contralateral sample, and contralateral test trials (colors as indicated), with shaded areas representing ±1 s.e.m. Panels show spatially selective ramp-down, midst, and ramp-up subtypes, as well as non-reward excitatory P-units, aligned to reward grasp (left) or forepaw contact (right). See Supplementary Table 1 for statistical results.

## Discussion

In this study, we investigated hippocampal oscillatory states and neuronal spikes during reward consumption in mice performing a DNMP T-maze spatial working memory task. We identified sparse reward-ensemble neurons in dorsal CA1 whose sustained firing was aligned to distinct phases of consummatory behavior and spatially centered on reward locations. These reward ensembles exhibited persistent firing coordinated with sensory-evoked ∼6-Hz theta oscillations and SWRs. Notably, TG-PAC was preferentially enhanced during sample trials, whereas SWR-triggered recruitment was elevated during test trials, particularly in spatially selective ramp-down and ramp-up units. This dissociation suggests that sensory encoding and post-choice associative processing are accompanied by distinct but interleaving oscillatory states. Atropine disrupted ∼6-Hz theta rhythmicity, gamma activity, and reward-unit firing, suggesting engagement of atropine-sensitive type 2 theta during consummation. These trial-dependent modulations were most prominent during the early phase of consummation, suggesting that sensory-associated cholinergic tone contributes to these dual oscillatory modes. Together, these findings indicate that the dual oscillatory mode during reward consumption may reflect the integration of ongoing sensory input into spatial working memory.

### Dual Oscillatory Modes During Consummatory Behavior

Unlike the classical classification of hippocampal oscillations based on behavior ^29^, we showed that reward consumption is accompanied by the dual oscillatory state of ∼6-Hz theta oscillations and SWRs during the early phase of consummation. Given that the hippocampus is crucial for linking initial sensory encoding to subsequent behavioral consequences, the dual oscillatory states in the reward units may provide a temporal window potentially favorable for memory processing during wakefulness. Previous studies have shown that awake SWRs contribute to working memory formation ^30,31^. A key prior observation is that brief pauses at reward locations boost SWR activation of CA1 pyramidal neurons with theta-mediated spatial fields centered at the reward locations ^27^, an effect that parallels NMDA-dependent spatial learning ^26^. In parallel, sensory input together with behavioral salience elevates SWR activity ^32,33^ and shapes its neuronal content ^34,35^. These findings suggest that the hippocampus integrates moment-to-moment sensory information with behavioral context. Nonetheless, previous work has neither demonstrated a direct link between theta and SWR states within an ongoing behavior, nor identified the specific CA1 neurons that may contribute to a spatiotemporal memory buffer for goal-directed behavior. We found that trial-dependent SWR enhancement was selectively observed at ipsilateral reward locations where spatially selective ramping units exhibited sustained theta modulation, whereas this modulation was absent at contralateral reward sites where their fields are weak. This pattern suggests coordinated engagement of theta and SWR neuronal dynamics at behaviorally salient locations during reward-guided memory processing. Together, the dual oscillatory state observed during reward consumption may support integration of ongoing sensory input with spatial working memory.

### Atropine-Sensitive Type 2 Theta as a Sensory–Linked State during Consummation

Atropine markedly disrupted both ∼6-Hz theta oscillations and reward-unit firing during consummation while sparing SWRs, suggesting engagement of atropine-sensitive type 2 theta during reward consumption ^36,37^. Type 2 theta is classically associated with immobility-linked sensory processing and cholinergic arousal ^15,16^. Accordingly, ∼6-Hz theta oscillations were phase-coherent with jaw movement rhythms during chewing, consistent with sensorimotor integration within theta ^38,39^. During consumption, theta power gradually decreased as SWR rates increased, although no clear temporal boundary separated the two oscillatory states. Instead, theta oscillations and SWRs coexisted during the early phase of consummation, with SWRs intermittently interrupting ongoing theta rhythms as previously reported ^27^. We also demonstrated that the reward-unit activity was significantly reduced upon arriving at empty reward locations during error trials, indicating dependence on sensory signals. In addition, the TG-PAC spike modulation was predominant at chewing onset, especially during sample trials, when encoding demands are expected to be highest. Reward units exhibited coherent spiking with both jaw and theta rhythms, whereas SWR peaks showed no phase locking to jaw movement rhythms, unlike nasal respiration that entrains awake SWRs ^40^. Several features of the observed oscillatory state are broadly consistent with proposed roles of cholinergic and supramammillary-septal circuits in coordinating sensory-linked theta activity during arousal and reward-related behavior ^41–43^. However, the present study does not directly resolve the circuit mechanisms underlying this coordination. Together, these findings suggest that atropine-sensitive type 2 theta may support integration of consummatory sensorimotor signals with hippocampal processing during reward-guided behavior.

### Putative circuit mechanisms underlying theta-SWR dual oscillatory modes

High-gamma oscillations in CA1 are classically linked to MEC III input, whereas low gamma has been associated with CA3 ^44,45^. High-gamma TG-PAC has been implicated in encoding-related processing ^46,47^, and low-gamma TG-PAC in retrieval ^48,49^. In contrast, we observed robust enhancement of low-gamma TG-PAC during the sample phase of reward consumption (Fig. 6). Recent studies have shown that low gamma oscillations can arise from CA2 ^50,51^, a region implicated in temporal integration ^52,53^. Consistently, theta slowed from ∼8 to ∼6 Hz after chewing onset, accompanied by a shift in reward-unit spiking from peak-phase firing to sustained locking on the descending phase and trough (Fig. 5), potentially reflecting shifts in theta-associated network coordination involving MEC III- and CA2-associated circuitry ^54^. Supporting a role in encoding, low-gamma TG-PAC strength scaled with spatial selectivity and was preferentially enhanced in spatially selective units during sample trials (Supplementary Fig. 13).

Features consistent with CA2-related modulation include persistent 6-Hz theta phase locking after chewing onset, elevated low-gamma TG-PAC, and spike timing preceding SWRs, all of which have been associated in previous studies with CA2 activity and pre-SWR recruitment ^30,55^. In parallel, a contribution from MEC III is suggested by elevated high-gamma TG-PAC, state-dependent theta phase shifts (Supplementary Fig. 6) ^25^, preferential peak-phase firing at reward grasp, strong spatial selectivity for reward locations, and localization to proximal CA1, the principal MEC III termination zone critical for spatial coding ^22,56–58^, a pathway in which ACh-dependent intrinsic persistent activity has been described ^59,60^. Together, these observations are consistent with the possibility that circuits associated with CA2 and MEC III contribute to theta-modulated spiking and transient integration of sensory and spatial information during consummatory behavior. However, the present study does not directly resolve the underlying circuit mechanisms.

Importantly, enhanced SWR activation during test trials was selectively observed in spatially selective ramping units exhibiting dual theta-SWR spiking modes (Figs. 7 and 8). This convergence of theta- and SWR-associated recruitment may facilitate integration of ongoing consummatory signals with spatiotemporal context and may promote plasticity linking reward consumption with spatial working memory representations ^61^.

Previous studies indicate that elevated cholinergic tone sustains theta oscillations and primes EC-HP circuits for encoding ^16,62–64^, whereas reduced cholinergic tone facilitates CA3–CA1 interactions required for retrieving and binding stored representations ^65–67^. Although this canonical framework often treats encoding- and retrieval-related states as temporally distinct, our findings suggest that these processes can occur concurrently through dual oscillatory states, potentially allowing ongoing sensory input to interact with existing memory representations. Interactions among CA2-, CA3-, and entorhinal-associated pathways may contribute to this coordination ^53,68,69^. However, the present study does not directly resolve how these circuits contribute to dual theta-SWR coordination in reward ensembles during consummatory behavior.

### Temporally structured reward ensembles

Our findings highlight that spatially selective reward location coding and non-selective coding are differentially distributed between ramp-down/up and midst units. A previous study utilizing two-photon calcium imaging combined with a virtual reality enclosure in a head-fixed condition demonstrated context-invariant reward location coding in ∼10% of CA1 neurons ^3^, a proportion akin to the reward units identified in our study. This discrepancy in the spatial selectivity of reward units may stem from variations in recording settings. We showed strong SWR-related spiking in reward units during consumption, which may obscure the spatial characteristics of the neuronal response. Furthermore, our DNMP T-maze task likely imposes greater spatial memory demands than their one-dimensional virtual reality task, where CA1 representations are particularly influenced by learning protocols employed ^70,71^. Moreover, differences in target regions may exist; our focus was primarily on proximal CA1, whereas Gauthier may have emphasized distal regions, including the Subiculum, where pyramidal neurons exhibit a greater number of firing fields ^72^, consistent with our midst units. Despite these differences, our study employed large-scale neural recordings from freely moving mice using custom-made 34-tetrode microdrives, which allowed us to capture hippocampal representations likely reflecting more natural sensory inputs. These recordings facilitated time-resolved analysis, including the correlation between neuronal oscillations and spikes, thereby enabling us to investigate critical hippocampal activity involved in spatial working memory. This organization is consistent with cognitive map framework ^2,7,73^.

We identified three distinct reward subtypes defined by their temporal coding during consummation. Unlike their diverse spatial selectivity, their temporal profiles remained largely invariant across trials and aligned to feeding events — grasping, chewing onset, and forepaw contact — rather than absolute time, distinguishing them from “time cells” ^74^. Ramp-down and ramp-up profiles resembled LEC III ramping signals that convey experience-based, trial-invariant temporal information ^75^. Ramp-down units initiated firing before reaching the reward or even an empty site during errors, whereas ramp-up units peaked at forepaw contact when theta power resumed. In contrast, midst units increased firing after chewing onset without a consistent peak, showed weak theta phase-locking, minimal REM phase shifting, and the lowest spatial selectivity. Along with their distal CA1 bias, these features are consistent with a greater influence of non-spatial variables and possibly from LEC III and proximal CA2/CA3 ^21,76,77^. A subset, however, adopted ramp-like profiles at contralateral reward sites, suggesting potential latent spatial learning roles. We also found a minority of reward units that consistently avoided SWR activation. These neurons may preferentially reflect internal sensory information at the reward and may be less engaged in contextual association. Given their low yield, whether these units represent preconfigured cell types or a reserve population for alternative contexts remains an open question.

### Broader implications for hippocampal-cortical coordination

More broadly, these dual oscillatory states may reflect state-dependent hippocampal-cortical coordination during attentive wakefulness. In this speculative framework, enhanced TG-PAC during sample trials could relate to online encoding, whereas enhanced SWR modulation during test trials could relate to post-choice integration of recent experiences. Such awake coordination may contribute to rapid updating of memory networks, distinct from sleep-associated systems consolidation ^78,79^.

## Acknowledgements

We thank all members of the Kitamura Laboratory for their support, Dr. Hisayuki Osanai for providing silicon probe recording data. This work was funded by Endowed Scholar Program (T.K) and Japan Society for the Promotion of Science grant 19J20331 (YO)

## Author Contributions

YO and TK conceptualized the study. YO, JY, and TK designed the experiments. YO, JY, and TK interpreted data. YO performed all experiments and analysis. YO designed the custom 34-tetrode microdrive, EIB, and preamplifier headstage used in this study. YO and TK wrote the manuscript, and JY provided comments on the manuscript.

## Data Availability

The data that support the findings of this study have been deposited in DANDI under accession number DANDI: 001775 and are publicly available as of the date of publication. Any additional information required to reanalyze the data reported in this paper is available from the corresponding authors upon reasonable request.

## Code Availability

The original code used for data analysis in this study has been deposited in ZENODO (DOI: 10.5281/zenodo.18580888) and is publicly available as of the date of publication.

## Methods

### Experimental Model and Subject Detail

All procedures involving mouse care and experimental manipulations were conducted in accordance with NIH and institutional guidelines and approved by the UT Southwestern Institutional Animal Care and Use Committee. Six male wild-type C57BL/6J mice (Jackson Laboratory; 6–12 months old) were used. Two mice recorded under different experimental conditions were excluded from the main cohort analyses unless otherwise noted: one implanted with an 18-tetrode microdrive and tested using flavored food rewards, and one implanted with a 32-channel silicon probe and recorded in the home cage. These animals were used primarily for atropine-related oscillatory analyses. Mice were pretrained to obtain food rewards on a raised T-maze track. Recording sessions were conducted after tetrodes stabilized in dorsal CA1. The main dataset consisted of recordings from five mice: A1 (12 tetrodes, 23 sessions), A2 (34 tetrodes, 15 sessions), A3 (34 tetrodes, 19 sessions), A4 (34 tetrodes, 15 sessions), and A5 (18 tetrodes, 7 sessions). At the conclusion of the experiments, brains were fixed, sectioned (40 μm), and tetrode tracks were verified histologically by gliosis-associated autofluorescence.

### Delayed Nonmatch-to-Place (DNMP) T-Maze Task

Food-deprived mice (maintained at 75% of pre-surgery body weight) were trained on a delayed nonmatch-to-place (DNMP) T-maze task (100 cm stem; 100 cm left and right goal arms) with food rewards (40–70 mg, Teklad Global 19% protein) placed at the ends of the left and right goal arms. Each trial consisted of a sample run followed by a test run separated by a 20 s delay, with 10–20 trials performed per daily session. During sample trials, one arm was blocked, forcing mice to retrieve reward from the opposite arm. During test trials, both arms were accessible, and reward was delivered only to the arm opposite the sample arm. Incorrect choices resulted in 120 s confinement at the non-rewarded site. Sample arms were pseudo-randomized to balance left and right trials. Mice were confined behind a guillotine door for 20 s before each run, and the maze was cleaned with 10% ethanol between trials. Sessions were preceded and followed by open-field exploration sessions (30 min pre-run; 30–120 min post-run). Behavioral position was tracked using two LEDs mounted on the microdrive headstage and recorded at 29.97 Hz using an overhead camera synchronized with Cheetah 6 software (Neuralynx).

### Electrode Implantation and Electrophysiology

Mice were anesthetized with isoflurane (0.5–2%) and implanted with independently movable tetrode microdrives targeting dorsal CA1 (AP −1.80 mm, ML ±1.60 mm, DV +2.20 mm). Implants consisted of 12 tetrodes (n = 1), 18 tetrodes (n = 1), or 34 tetrodes (n = 3). In the 34-tetrode configuration, 17 tetrodes were implanted in each hemisphere, whereas 12- and 18-tetrode implants targeted the right hemisphere only. A separate mouse used for atropine-related home-cage recordings was implanted with a 32-channel linear silicon probe. Signals were acquired using Neuralynx acquisition systems at 32 kHz with skull-screw grounding and corpus callosum references. Microdrives were designed in SolidWorks and 3D-printed in resin. After four days of recovery, tetrodes were gradually advanced toward CA1 over 7–14 days while monitoring sharp-wave ripple (SWR) activity, with advances limited to <40 μm/day near the pyramidal layer. Tetrodes were targeted to the deep pyramidal layer based on SWR polarity ^25^. Recordings began after tetrode stabilization and behavioral training (>70% correct choice). Unit activity was band-pass filtered at 600–8,000 Hz and local field potentials (LFPs) at 1–500 Hz. Offline spike sorting was performed using MountainSort4 112 followed by manual curation using custom software.

### Definition of Feeding Behavioral Sequence

Feeding behavior was recorded using side-view cameras positioned 15 cm from each reward well and synchronized with neural activity via Cheetah software. Mice exhibited a stereotyped feeding sequence consisting of reward grasp, chewing while holding the pellet with the forepaws, and forepaw placement on the floor following consumption. The consummatory epoch was defined as the interval between reward grasp and forepaw placement. Three behavioral timestamps were annotated: 1) reward grasp, 2) chewing onset, and 3) forepaw placement. Timestamps were generated by TTL triggering through a switch connected to the Neuralynx system and subsequently verified by video inspection using custom software. Trials in which mice struggled to collect food or dropped pellets during consumption were excluded from the main analyses.

### Classification of Excitatory and Inhibitory Units

Unit classification was performed following Kay et al. ^52^. Well-isolated units (isolation metric >0.95, noise overlap <0.03, and cluster signal-to-noise ratio >2) were classified based on average firing rate, spike width, and the mean spike autocorrelation measure (ACM) computed over the 0–50 ms range using 0.1 ms bins. Units were classified as inhibitory if they exhibited short spike widths, ACM >25 ms, or average firing rates >10 Hz. Units not meeting these criteria were classified as excitatory. Classification was further verified visually using spike feature raster plots.

### Identification of Reward Units with Temporal Patterns

Reward units were analyzed separately for left- and right-arm trials. Inhibitory units and excitatory units with firing rates <1 Hz during consummation, excluding spikes within SWRs, were excluded. Instantaneous firing rates (IFRs) were computed from reward grasp to forepaw placement by convolving spike trains binned at 1 s with a Gaussian kernel (σ = 1.5 s), followed by duration normalization. IFRs were averaged independently for left- and right-arm trials. Significant temporal modulation was assessed across 20 normalized time bins using shuffle-based ANOVA with 1,000 circular spike-time shuffles within the consummatory epoch; units exceeding the 95th percentile of the shuffled distribution were retained. Population dynamics were quantified using spike occurrence ratios across seven time bins, with the first and last bins fixed at 5 s and the intermediate bins equally spaced and normalized to 5 s duration. Spike counts were normalized to total spikes within each unit. Temporal profiles were classified using principal component analysis followed by agglomerative hierarchical clustering with average linkage and cosine distance. The clustering threshold was defined midway between the third-from-last and second-from-last linkage distances.

### Reward Location Spatial Selectivity and Sample-Test Trial Selectivity

Spatial selectivity index (SI) for each reward unit was calculated as follows. For choice selectivity, SI was defined as (*FR_R_* − *FR*_*L*_)/(*FR_R_* + *FR*_*L*_) for right trials and (*FR*_*L*_ − *FR*_*R*_)/(*FR*_*R*_ + *FR*_*L*_) for left trials, where *FR*_*R*_ and *FR*_*L*_ are trial-averaged firing rates during reward consumption computed from spikes outside SWRs. For sample-test trial selectivity, SI was defined as (*FR*_*S*_− *FR*_*T*_)/(*FR*_*S*_+ *FR*_*T*_), where *FR*_*S*_ and *FR*_*T*_ are trial-averaged firing rates from spikes outside SWRs in sample and correct-choice test trials, respectively. The significance of the selectivity indices was assessed using a two-way ANOVA with factors of choice and trial type (p < 0.05), defining spatially selective and trial selective reward units.

### Comparison of Individual Reward Unit Subclasses Between Choice Arms

To assess whether reward unit subclasses were preserved across choice arms, subtype identities assigned during left- and right-arm trials were compared for each unit. A null distribution was generated from 5,000 shuffled subtype assignments under the assumption that subtype identity was independent between arms. Observed subtype pairings were converted to z-scores relative to the shuffled distribution, such that positive z-scores indicated subtype combinations occurring more frequently than expected by chance, whereas negative z-scores indicated combinations occurring less frequently than expected.

### SWR Detection and LFP Spike Modulation

Sharp-wave ripples (SWRs) were detected from local field potential (LFP) recordings in the CA1 pyramidal layer. SWRs were defined as transient events in which the square-rooted ripple-band signal (120–250 Hz), smoothed with a Gaussian kernel (σ = 4 ms), exceeded 3 standard deviations (SD) above the mean of each tetrode. SWR epochs were defined from the ripple peak to the time points at which ripple-band power returned below 1 SD. Neighboring events were merged if separated by <20 ms or if their 1 SD boundaries were <10 ms apart. Events shorter than 15 ms were excluded. Spikes occurring within SWR epochs were classified as SWR spikes. Spikes occurring at least ±250 ms from SWR peaks were classified as non-SWR spikes and used for subsequent analyses, including pairwise phase consistency (PPC2), theta-gamma phase-amplitude coupling (TG-PAC), spike theta-phase autocorrelograms, and spike jaw-phase locking. Only tetrodes localized to the CA1 pyramidal layer based on SWR polarity and spike waveform characteristics were included in theta-phase analyses.

**Pairwise Phase Consistency (PPC2)** ^28,80^ was used to quantify spike-LFP phase coupling from continuous wavelet spectrograms computed from CA1 LFPs using the Amor analytic wavelet. PPC2 was calculated for each frequency during reward consumption using 2.5 s sliding windows with 1.25 s steps. PPC2 values were computed across spike-LFP phase pairs from separate trials and normalized by the total number of spike combinations. PPC2 values were z-scored relative to 100 within-window circular spike-time shuffles and averaged across tetrodes. Time-resolved peak synchronization frequencies were calculated from the frequency exhibiting maximal PPC2 within each time bin. To assess relationships between spatial selectivity and theta synchronization, Kendall’s tau correlations were computed between spatial selectivity indices and PPC2 values averaged across the 4–10 Hz theta band. Statistical significance was evaluated using 5,000 within-window circular spike-time shuffles. Circular-linear correlations between preferred theta phase and spatial selectivity were computed for each reward subtype and assessed using permutation testing. Differences in synchronization frequency between spatially selective and non-selective units were evaluated using cluster-based permutation tests ^81^. Cluster-level statistics were defined as the summed t-values of contiguous supra-threshold clusters (500 permutations; two-sided p < 0.05).

**Spike Theta-Phase Autocorrelograms** ^22^ were computed from the same wavelet spectrograms to quantify spike rhythmicity relative to ongoing theta oscillations (3–12 Hz). Theta phase at each time point was obtained by circularly averaging phase values across the theta band, and spike phases were assigned by linear interpolation. Phase differences between spikes were computed using 2.5 s sliding windows with 1.25 s steps and 30° phase bins, then averaged across trials. Autocorrelograms were z-scored relative to 500 within-window circular spike-time shuffles and subsequently averaged across recording channels.

**Theta-Phase Gamma-Power Coupling (TG-PAC)** ^82^ was quantified at spike times to assess cross-frequency coupling between theta phase (3–12 Hz) and gamma amplitude in low-gamma (30–55 Hz) and high-gamma (60–120 Hz) bands. For each spike, phase-amplitude coupling (PAC) was measured within ±600 ms using 5 s sliding windows with 2.5 s steps during reward consumption. Peri-spike PAC values were z-scored relative to 100 within-window circular spike-time shuffles. For each unit, TG-PAC values were averaged across theta phase, gamma power, and spike time lag within ±10 ms separately for sample and test trials. Trial-dependent TG-PAC modulation across reward subtypes was evaluated using paired t-tests between sample and test trials. Only units exhibiting TG-PAC values with Z ≥ 1 in both trial types were included. To determine whether modulation occurred specifically at spike timing, peri-spike TG-PAC differences between sample and test trials were further evaluated using cluster-based paired permutation tests. Because the directional hypothesis predicted greater TG-PAC during sample than test trials, significance was assessed using one-sided tests (500 shuffles, p < 0.05). Differences in peri-spike TG-PAC timing between spatially selective and non-selective units were assessed similarly using cluster-based permutation tests. Relationships between spatial selectivity and TG-PAC modulation were evaluated using Kendall’s tau correlations computed in 2.5 s sliding windows with 1.25 s steps. Statistical significance was assessed using null distributions generated from 2,000 random permutations of TG-PAC values across units.

**SWR Spike Modulation** was quantified using peri-SWR spike histograms constructed with 7 ms bins over a ±300 ms window around SWR peaks. For time-resolved analyses (Figs. 7 and 8), histograms were computed using 5 s sliding windows with 2.5 s steps and z-scored relative to 500 within-window circular spike-time shuffles generated separately for each window. For trial-averaged analyses (Fig. 3; Supplementary Figs. 14 and 15), histograms were computed across the entire consummatory period. SWR spike modulation was quantified as the mean z-scored peri-SWR spike rate within the −30 to 0 ms window relative to SWR peaks. Trial-dependent modulation of SWR-related spiking was assessed using one-sided paired t-tests (test > sample). Differences in peri-SWR spike histograms between trial types were further evaluated using cluster-based paired permutation tests analogous to the TG-PAC analysis. Because the directional hypothesis predicted greater SWR-related firing during test than sample trials, significance was assessed using one-sided tests (500 shuffles, p < 0.05).

**SWR-Positive and SWR-Negative Units** were classified based on SWR-associated activity. Units were classified as SWR-positive if the mean z-score within ±21 ms of SWR peaks exceeded the mean z-score outside this window, and as SWR-negative if the mean z-score within ±21 ms was lower than that outside the window. All spikes occurring within SWR events during reward consumption were included in this analysis.

**REM Theta-Phase Preference** was quantified using the Hilbert transform of band-pass-filtered LFPs (4–12 Hz; FIR filter, order 2,000). For each unit, theta phases at spike times were circularly averaged to determine preferred theta phase. As a control, theta phase during RUN epochs in the open field was computed similarly to compare preferred theta phase distributions between REM sleep and active exploration. REM epoch detection is described below.

### Chewing Rhythms Spike Modulation

Jaw movements were recorded using front-view cameras capturing the maxilla and mandible. Body-part positions were tracked using DeepLabCut (DLC) ^83^ and included only when likelihood scores exceeded 0.95 for at least 1 s. Inter-jaw distance was computed and used for spectrogram and spectral analyses of chewing rhythmicity. Power spectra of jaw displacement signals (1–14 Hz) were computed using Welch’s method with a 512-sample Hamming window, 75% overlap, and 1024 discrete Fourier transform points. Jaw phase at spikes occurring both within and outside SWRs was estimated using the Hilbert transform of band-pass-filtered jaw displacement signals (FIR filter, order 1,000) with linear interpolation. Theta phase at spike times was computed similarly from CA1 pyramidal layer LFPs filtered at 3–12 Hz. Jaw- and theta-phase locking during consummation were quantified using weighted circular concentration statistics corrected for non-uniform phase occupancy (20 phase bins spanning [−π, π]). Statistical significance was assessed using null distributions generated from 2,000 circular spike-time shuffles. Spike co-modulation between jaw and theta rhythms was assessed using circular-circular correlation. Relationships between spatial selectivity and jaw-rhythm synchronization were evaluated using circular-linear correlations between preferred jaw phase and spatial selectivity indices, as well as Kendall’s tau correlations between Rayleigh’s Z scores and spatial selectivity indices. Statistical significance was assessed using permutation testing with 2,000 shuffled spike trains.

### Atropine Injection and LFP Analysis

Three mice were used to assess the effects of atropine sulfate on CA1 local field potential (LFP) oscillations. Saline (0.9%, volume matched) or atropine sulfate (50 mg/kg, i.p.) was administered 15 min before recordings. Two mice implanted with tetrodes performed the DNMP T-maze task under saline and atropine conditions, whereas a third mouse implanted with a 32-channel linear silicon probe was recorded during voluntary food consumption in the home cage to examine depth-resolved CA1 oscillatory activity. Food consumption was monitored by video recording, and uninterrupted 12 s epochs following chewing onset were used for spectral analyses. Power spectral densities were computed using Welch’s method with a 512-sample Hamming window, 75% overlap, and 1024 discrete Fourier transform points. Spectral periodicity was quantified using the Fitting Oscillations and One Over F (FOOOF) model ^84^ to separate periodic and aperiodic spectral components. Aperiodic fitting ranges were set to 1–30 Hz for theta, 20–120 Hz for gamma, and 100–249 Hz for ripple-band analyses. Periodic power was quantified from residual spectra averaged over 3–9 Hz (theta), 35–80 Hz (gamma), and 120–249 Hz (SWR). Spectral power values were pooled across trials and compared between saline and atropine conditions using unpaired t-tests. To assess synchronization between recording sites in the CA1 pyramidal layer during reward consumption, magnitude-squared coherence was computed between LFP pairs. Coherence values ranged from 0 (no correlation) to 1 (perfect correlation).

### Detection of REM Sleep

Three mice were used to investigate theta-spike modulation during REM sleep. REM epochs were initially identified online by monitoring immobility (<2 cm/s) together with prominent theta oscillations across recording channels and marked using TTL-triggered signals. Epoch boundaries were subsequently refined manually using custom software. Theta (4–12 Hz) and delta (1–4 Hz) power, together with spectrograms, were inspected to verify behavioral state transitions. A total of 682 s of REM sleep was included in the analyses. As a control condition, RUN epochs were defined during open-field exploration before T-maze trials when head velocity exceeded 4 cm/s, yielding a total of 1,237 s of RUN epochs.

### Occupancy-Normalized Firing Map

For each unit, occupancy-normalized firing rate maps were computed from spikes occurring during T-maze run epochs, excluding periods when mice were transferred from the reward site to the start position. Spike counts were divided by occupancy time within 2 cm × 2 cm spatial bins and smoothed using a two-dimensional Gaussian kernel (σ = 3 cm). Peak firing rate was defined as the maximum value of the resulting firing rate map.

### Supplemental silicon probe recording and ICA-based EMG extraction

A single-shank 32-channel silicon probe (50 μm site spacing; E32+R-50-S1-L6 NT, ATLAS Neuroengineering, Leuven, Belgium) mounted on a microdrive was implanted into dorsal CA1 in an isoflurane-anesthetized 8-month-old mouse (AP, -1.80 mm; ML, +1.65 mm; DV, +2.20 mm). A cerebellar skull screw served as ground and reference. Signals were acquired with Open Ephys at 20 kHz, and LFPs were obtained by band-pass filtering (0.1-500 Hz) and downsampling to 500 Hz. EMG-related activity was extracted from multichannel LFPs using independent component analysis (ICA), as previously described ^85^.

### Statistics

All analyses were performed using custom MATLAB R2019b scripts unless otherwise stated. Population averages are reported as mean ± standard error of the mean (SEM). Circular statistics were performed using CircStats ^86^. Unless otherwise specified, statistical significance was defined as p < 0.05. Post hoc comparisons following analysis of variance (ANOVA) were corrected for multiple comparisons. Statistical analyses were primarily performed at the single-unit level because reward-ensemble dynamics were defined based on individual neuronal activity, consistent with previous hippocampal electrophysiology studies. Cross-animal consistency was additionally evaluated where appropriate. For LFP-spike modulation analyses, including SWR spike modulation and TG-PAC modulation, comparisons between sample and test trials were performed using balanced trial counts across conditions.

