## Supplementary Information for "Hippocampal consummatory reward ensembles dynamically engage theta-SWR states during spatial working memory"

Title

Author Information

**Authors:**
Yoshiyuki Omura[1,4]*, Jun Yamamoto[1-3]* & Takashi Kitamura[1-3]*

**Affiliations**:
[1]Department of Psychiatry, University of Texas Southwestern Medical Center, Dallas, TX 75390, USA
[2]Department of Neuroscience, University of Texas Southwestern Medical Center, Dallas, TX 75390, USA
[3]Peter O'Donnell Brain Institute, University of Texas Southwestern Medical Center, Dallas, TX 75390, USA
[4]Lead contact
*Corresponding authors:
,



### Supplementary Figures

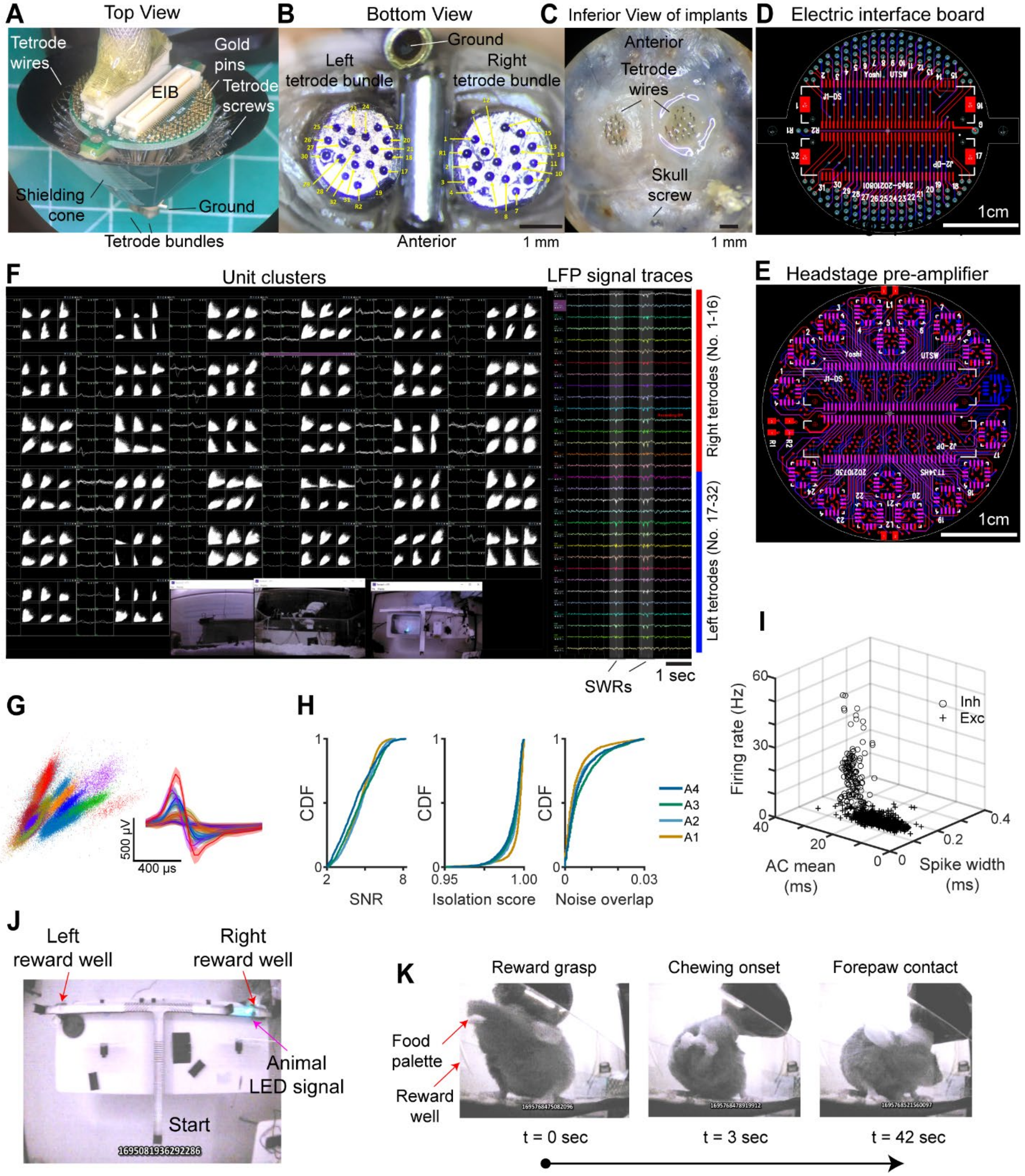

Supplementary Figure 1. Large-scale neural recording and behavioral tracking platform for freely moving mice (related to Figure 1).

**(A)** Top view of the fully loaded 34-tetrode microdrive showing tetrodes connected to the custom electrical interface board (EIB). **(B)** Bottom view of the microdrive showing tetrode tips and their spatial distribution targeting bilateral CA1 regions. **(C)** Inferior view of the implant following brain extraction, showing protruding tetrodes and their arrangement relative to the skull. **(D)** Layout of the custom-designed EIB used for tetrode signal routing. **(E)** Layout of the custom headstage pre-amplifier showing connections to the EIB and operational amplifier circuitry used for signal amplification prior to acquisition. **(F)** Representative neuronal spike clusters and LFP traces recorded simultaneously across tetrodes, demonstrating successful multi-unit isolation (234 units total) and signal quality. Negative polarity of sharp-wave ripples (SWRs) across tetrode channels indicates positioning within the deep CA1 pyramidal layer. **(G)** Left, representative spike clusters plotted in two-dimensional feature space using peak amplitudes from channels 1 and 2. Spikes were sorted using MountainSort4. Right, corresponding spike waveforms recorded from channel 1 (mean ± 1 s.d.). **(H)** Cumulative distributions of signal-to-noise ratio (SNR), isolation score, and noise-overlap values for isolated units across animals (A1, 12 tetrodes; A2–A4, 34 tetrodes). **(I)** Three-dimensional spike-feature distributions based on spike width, mean autocorrelation, and firing rate. Excitatory and inhibitory units were separated based on waveform and firing properties. **(J)** Top-camera behavioral tracking during the DNMP T-maze task. The headstage LED was tracked at 29.97 Hz using Cheetah software (Neuralynx). Example frame shows reward consumption at the right reward well. A guillotine door was positioned behind the animal during consummation. **(K)** Side-camera views of behavioral events defining the consummatory period: reward grasp (left), chewing onset (middle), and forepaw contact (right). The consummatory epoch was defined as the interval between reward grasp and forepaw contact.

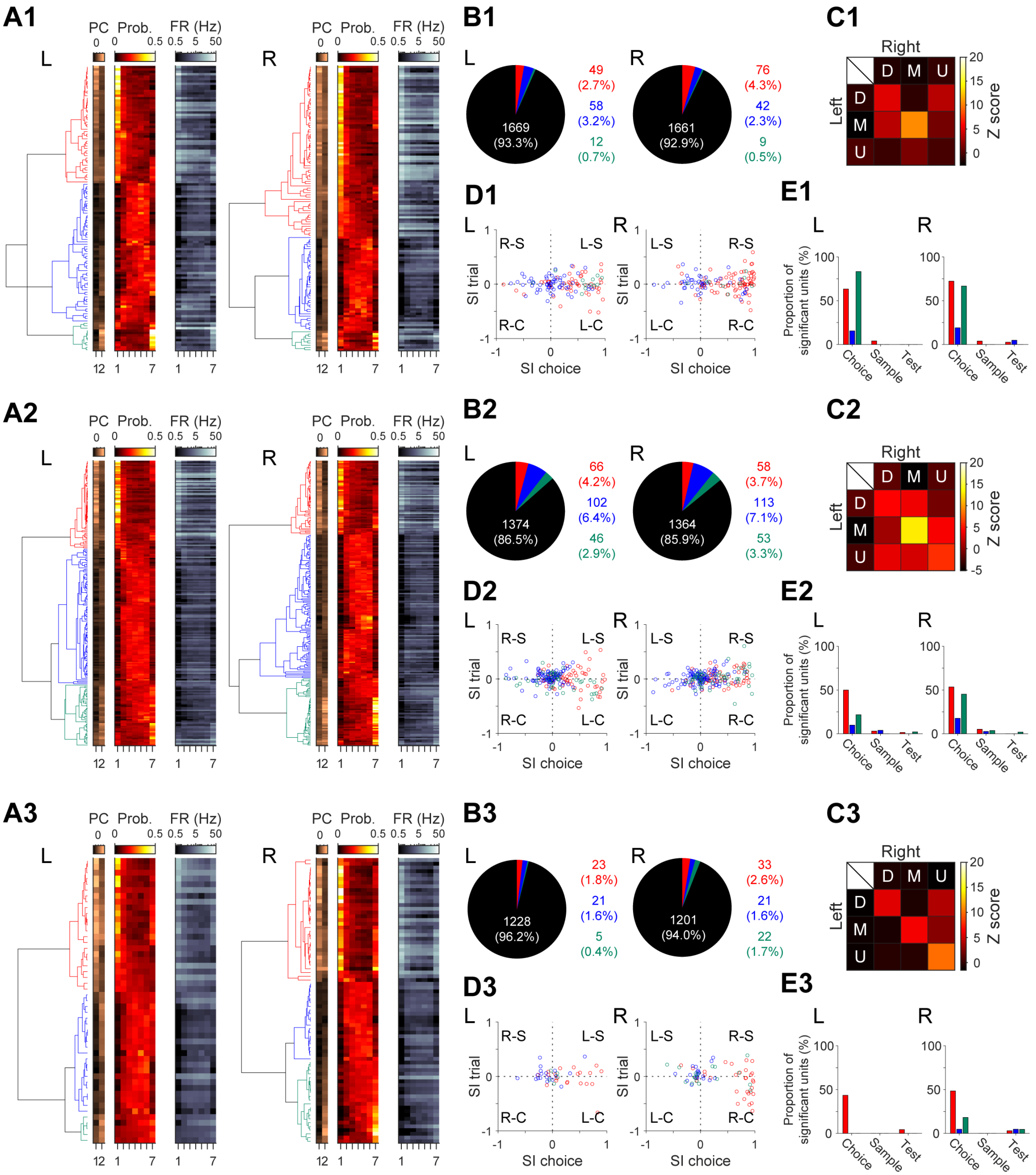

Supplementary Figure 2. Reward-unit classification across individual animals (related to Figure 2). From top to bottom, data are shown for animals A1 (12-tetrodes), A3 (34-tetrodes), and A4 (34-tetrodes). Data from A2 are shown in Figure 2. A5 was excluded from the main cohort analyses as described in Methods. Same analyses and plotting conventions as in Figure 2. See Supplementary Table 1 for statistical results.

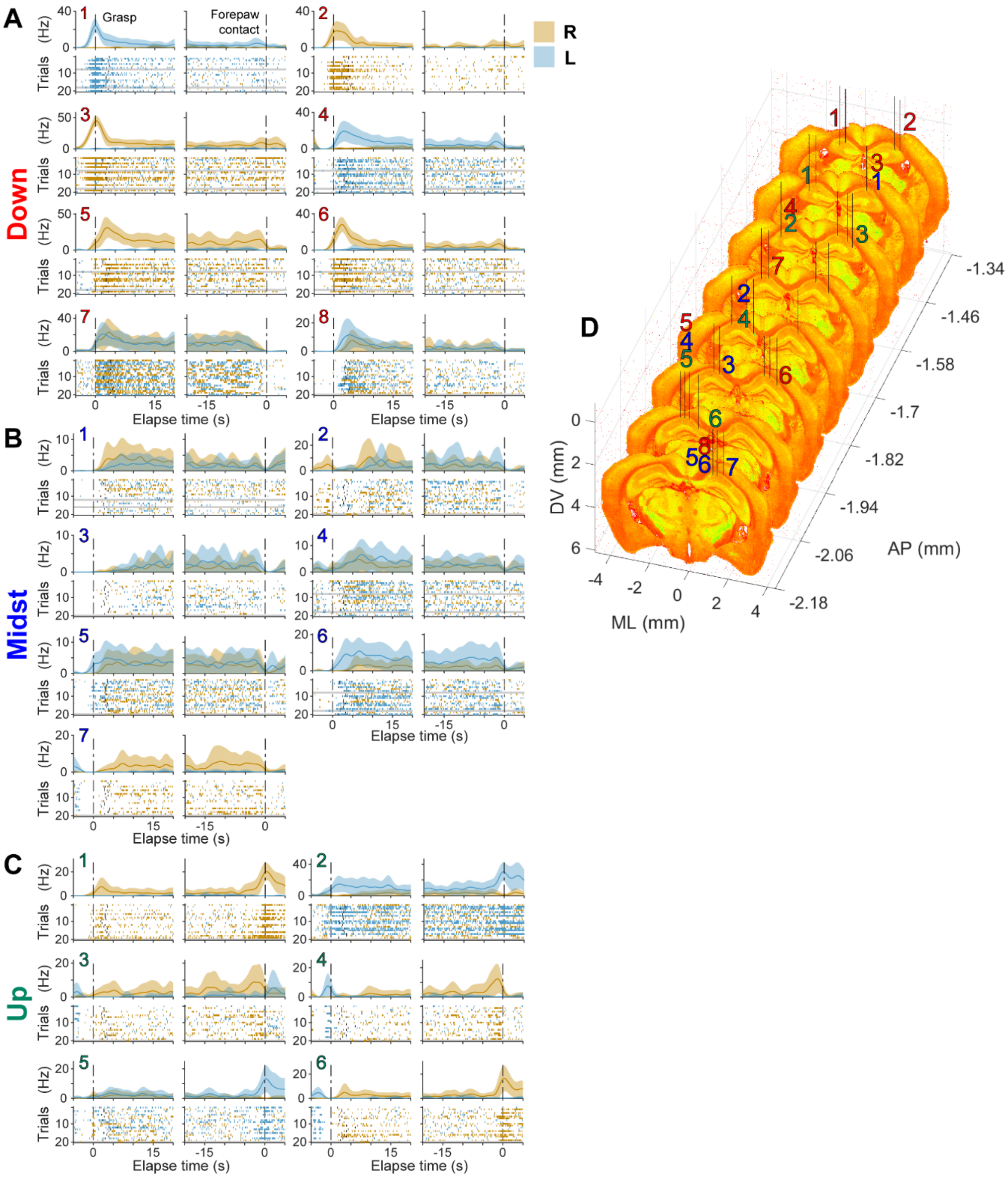

Supplementary Figure 3. Representative firing patterns of reward-unit subtypes (related to Figures 1 and 2).

Left, spike rasters and trial-averaged firing rates for representative ramp-down (red), midst (blue), and ramp-up (green) units shown separately for right- and left-arm trials. Black vertical dots indicate chewing onset, and gray shaded regions denote error trials. Right, histological reconstruction of tetrode locations from animal A2. Colored numbers indicate units corresponding to the spike rasters and firing-rate plots shown on the left. AP, anterior-posterior; ML, mediolateral; DV, dorsoventral coordinates.

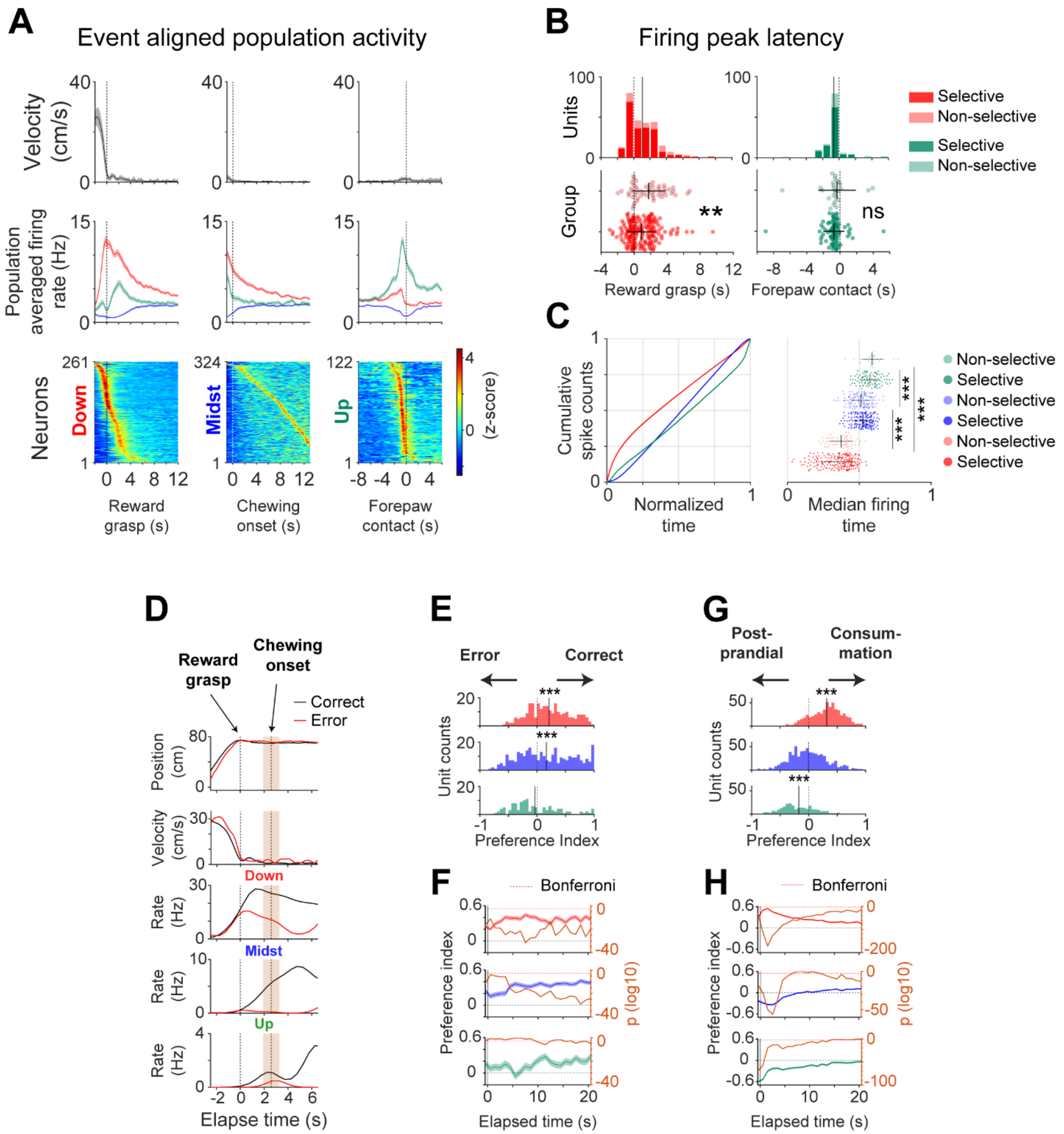

**Supplementary Figure 4. Population firing dynamics across consummatory, error, and postprandial epochs (related to Figure 2).**

**(A)** Population-averaged firing activity aligned to reward grasp, chewing onset, and forepaw contact. From top to bottom: head velocity, population-averaged firing rate (mean ±1 s.e.m.), and z-scored firing-rate heatmaps for all reward units. Right- and left-arm units were pooled. **(B)** Distribution of peak firing latencies relative to reward grasp (left) and forepaw contact (right). Bottom, each dot represents one unit. Spatially selective and non-selective units are shown in different color tones. **(C)** Population-averaged cumulative spike-count distributions for each subtype. Left, consummatory epochs normalized from 0 (reward grasp) to 1 (forepaw contact), with spike counts normalized from 0 to 1. Right, raster plots showing median firing times for individual units, defined as the time at which 50% of spikes occurred. ***$p < 0.001$. **(D)** Representative firing activity during correct and error trials. From top to bottom: head position, head velocity, and firing rates of representative ramp-down, midst, and ramp-up units. **(E)** Preference index comparing firing rates between correct and error trials during the 20 s period following reward grasp or arrival at an empty reward well. Preference indices ranged from −1 (error-biased) to 1 (correct-biased). **(F)** Population-averaged time-resolved preference indices (left axes) and corresponding p values (right axes) comparing firing rates between correct and error trials. Red dashed lines indicate the Bonferroni-corrected significance threshold. **(G)** Preference index comparing firing rates between consummatory and postprandial immobility epochs during the 20 s period following reward grasp or forepaw contact. **(H)** Population-averaged time-resolved preference indices comparing firing rates between consummatory and postprandial immobility epochs aligned to reward grasp and forepaw contact. See Supplementary Table 1 for statistical results.

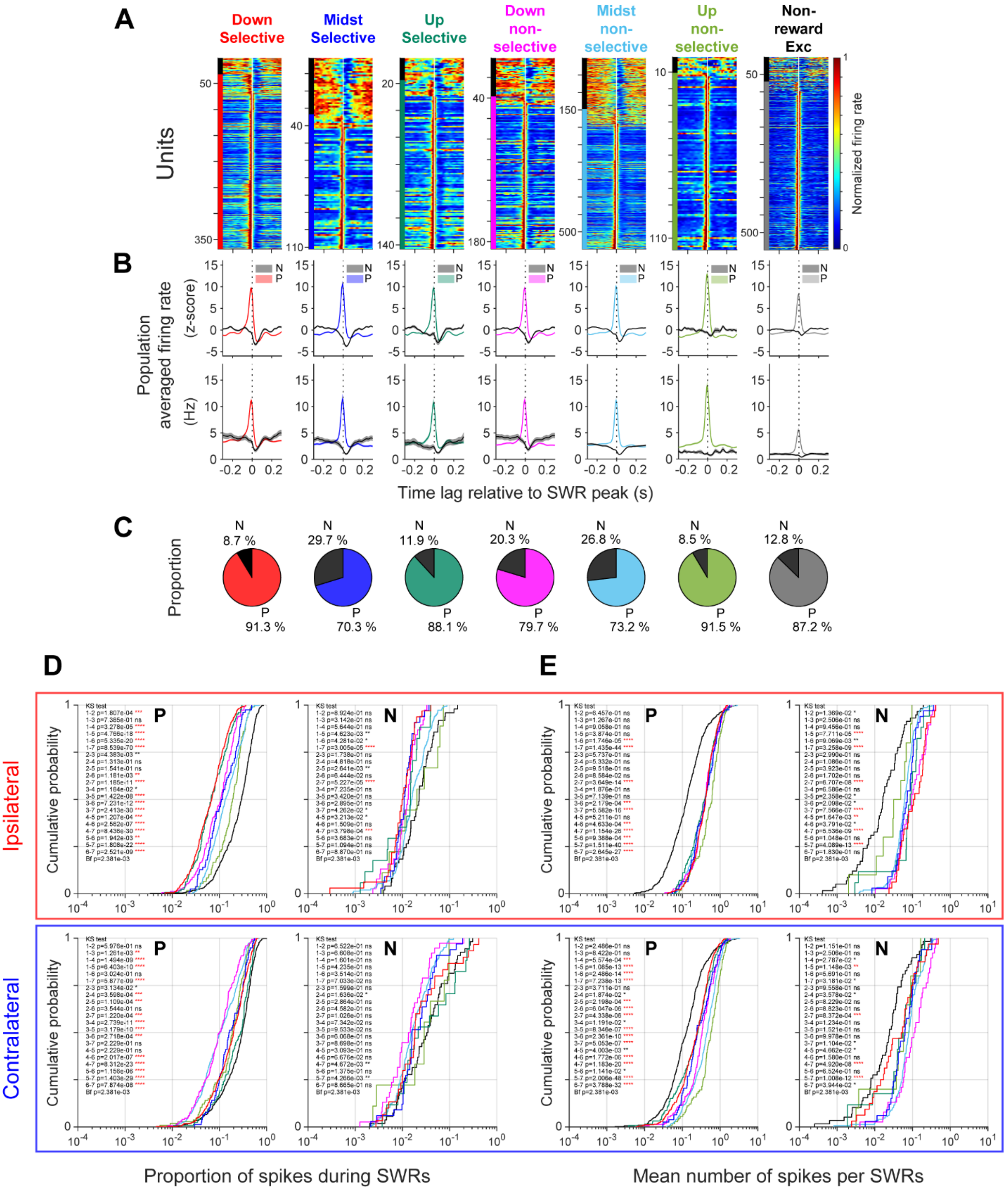

Supplementary Figure 5. Reward-unit activity during SWRs in reward consummation (related to Figure 3).

**(A)** Peri-SWR firing histograms of spatially selective reward, non-selective reward, and non-reward excitatory units aligned to SWR peaks (white vertical line). Firing rates were normalized to the maximum firing rate of each unit. Left color bars indicate P-units and N-units (black). **(B)**

Population-averaged z-scored peri-SWR firing histograms and mean firing rates (Hz) across reward subtypes and non-reward excitatory units. **(C)** Proportions of P-units and N-units for spatially selective reward subtypes, non-selective reward subtypes, and non-reward excitatory units. **(D)** Cumulative distributions of the proportion of spikes occurring during SWRs for individual units during the consummatory period. Colored lines indicate reward subtypes and non-reward excitatory units, with P-units and N-units plotted separately. Top, ipsilateral reward arm corresponding to the choice arm in which each unit was defined. Spatially selective ramp-down and ramp-up units exhibited lower proportions of SWR-triggered spikes than non-reward excitatory units, indicating sustained firing outside SWRs. Inset numbers indicate unit classes: 1, spatially selective ramp-down; 2, spatially selective midst; 3, spatially selective ramp-up; 4, spatially non-selective ramp-down; 5, spatially non-selective midst; 6, spatially non-selective ramp-up; 7, non-reward excitatory units. Statistical differences between cumulative distributions were assessed using the Kolmogorov-Smirnov (KS) test (*$p < 0.05$, **$p < 0.01$, ***$p < 0.001$, ****$p < 0.0001$). Red asterisks indicate significance after Bonferroni correction. Bottom, proportion of SWR-triggered spikes at the contralateral reward arm for the same units. Spatially selective ramp-down and ramp-up units exhibited rightward-shifted distributions at contralateral reward sites (related to Figure 8). **(E)** Same as in D, but cumulative distributions show the mean number of spikes per SWR for individual units.

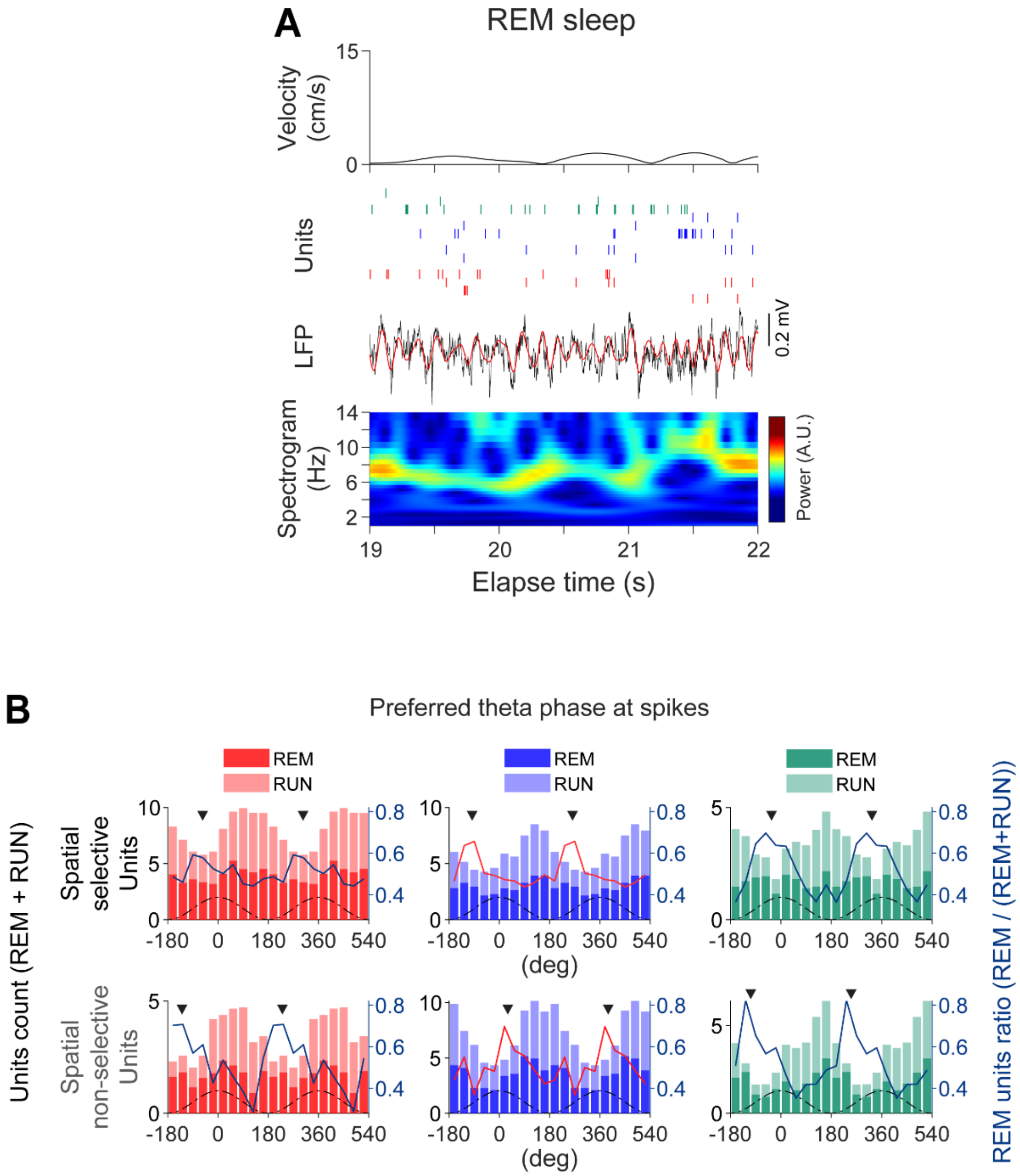

Supplementary Figure 6. Reward-unit activity during REM sleep (related to Figure 3).

**(A)** Representative theta-rhythmic activity during REM. From top to bottom: head velocity, spike rasters of representative units, hippocampal LFP trace, and LFP spectrogram showing theta oscillations during REM sleep. **(B)** Distributions of preferred theta phases for reward units during REM sleep and running (RUN) plotted across two theta cycles. Stacked histograms show phase preferences for units exhibiting significant phase locking in both behavioral states (Rayleigh test, $p < 0.05$). Overlaid lines indicate the ratio of units preferring each theta phase during REM sleep relative to RUN.

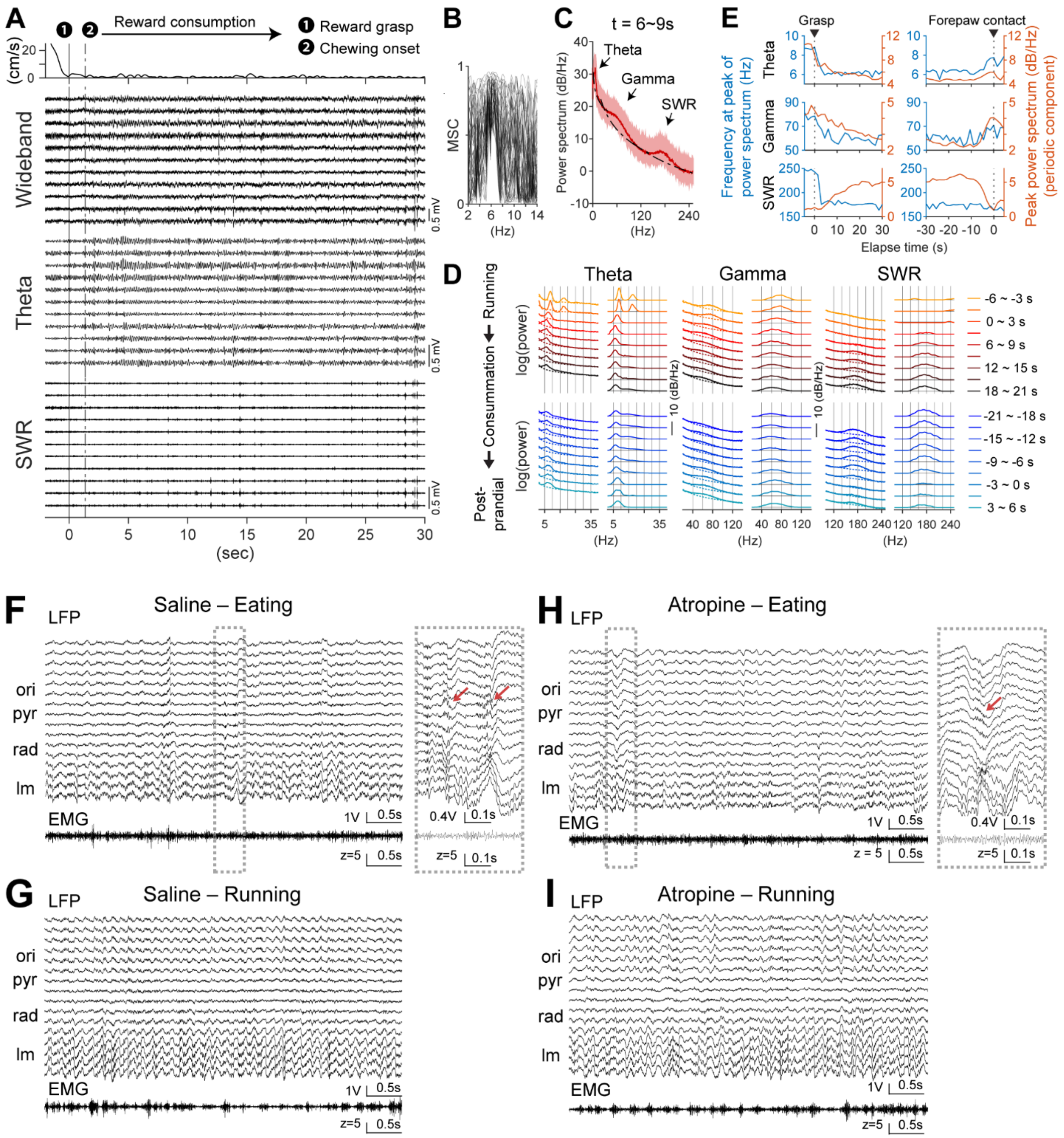

Supplementary Figure 7. Spectral characterization of 6-Hz oscillations during consummation and their sensitivity to atropine (related to Figure 4).

**(A)** From top to bottom: head velocity aligned to reward grasp; simultaneously recorded wideband hippocampal LFP traces from 11 channels in right dorsal CA1; theta-band (3–12 Hz) band-pass-filtered LFP traces from the same channels; and SWR-band (120–250 Hz) filtered LFP traces. Theta oscillations and SWRs occurred within the same consummatory period. **(B)** Magnitude-squared coherence computed across all pairwise combinations of the 11 CA1 LFP channels shown in A. Coherence values range from 0 (no correlation) to 1 (perfect correlation).

High coherence in the theta range indicates synchronized theta oscillations across CA1 channels during consummatory immobility. **(C)** Power spectral density (PSD) of CA1 LFP computed using Welch's method in 3 s time bins during reward consumption. Spectral periodicity was quantified using the FOOOF model (Donoghue et al., 2020) to separate periodic and aperiodic spectral components. PSDs show mean power spectra with 95% confidence intervals during mid-consummation (6–9 s after reward grasp). **(D)** Time-resolved PSD analysis aligned to reward grasp (top) and forepaw contact (bottom). Corresponding right panels show periodic spectral components obtained after subtraction of fitted 1/f components. Theta–gamma and SWR oscillations occurred during the initial phase of consummation. **(E)** Time-resolved peak frequencies (left axes) and corresponding peak periodic power (right axes) for theta, gamma, and SWR oscillations. Left panels are aligned to reward grasp; right panels are aligned to forepaw contact. **(F)** Noise-suppressed LFPs (top; ICA-based removal of EMG-related noise) and EMG traces (bottom) during consummatory behavior recorded 15 min after saline intraperitoneal injection using a 12-channel silicon probe in right dorsal CA1. Red arrow indicates a SWR event. **(G)** Noise-suppressed LFPs and EMG traces during running in an open field recorded 15 min after saline injection. Theta oscillations during running were faster than those during consummation. **(H)** Noise-suppressed LFPs and EMG traces during consummation recorded 15 min after atropine intraperitoneal injection. Red arrow indicates a SWR event. Theta rhythmicity was reduced relative to saline. **(I)** Noise-suppressed LFPs and EMG traces during running recorded 15 min after atropine injection. Theta rhythmicity remained largely preserved.

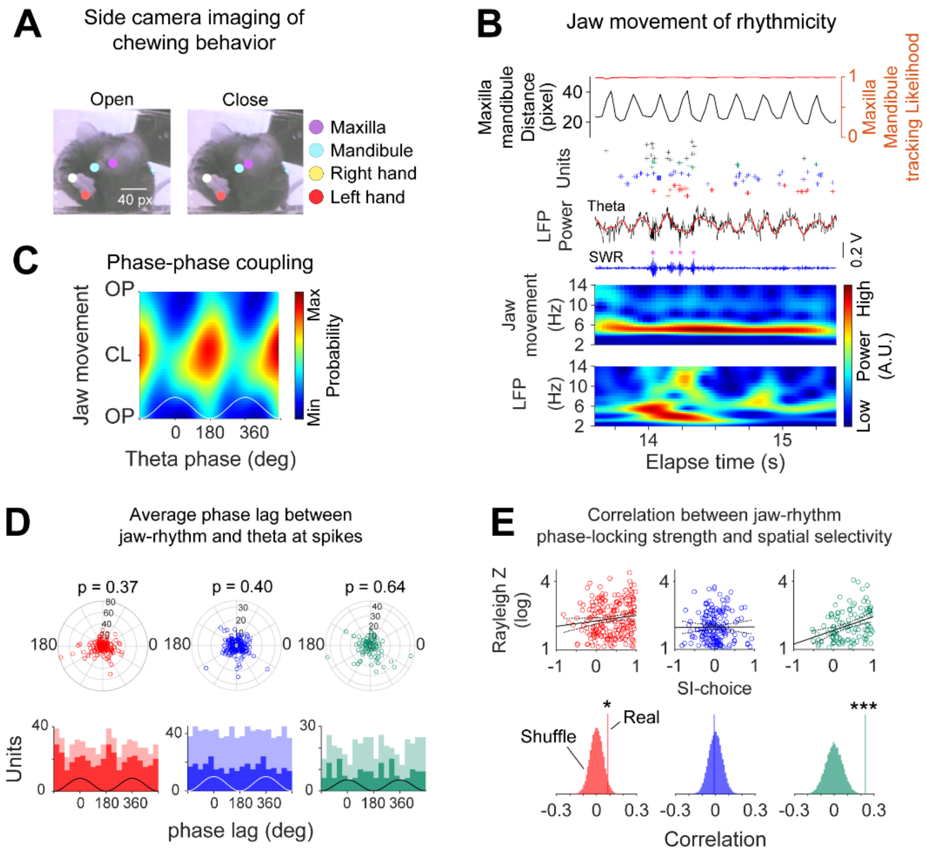

Supplementary Figure 8. Phase synchronization between theta oscillations and chewing rhythms (related to Figure 4).

**(A)** Side-mounted camera tracking of jaw movement rhythmicity during reward consumption. Maxilla and mandible movements were tracked using DeepLabCut. **(B)** Jaw movement rhythmicity, reward-unit activity, and hippocampal LFP traces during consummatory behavior. Top, jaw movement rhythmicity defined by the distance between the maxilla and mandible using body-part tracking likelihoods >0.95. Bottom, spike rasters of representative reward units, theta- and SWR-band filtered LFP traces, and wavelet-transformed jaw movement rhythmicity and hippocampal LFP signals. Both rhythms exhibited ~6-Hz periodicity. **(C)** Phase-phase coupling heatmap between jaw movement rhythmicity and theta oscillations. Heatmap shows the probability density of phase relationships between the two rhythms. OP, mouth opening; CL, mouth closing. **(D)** Distribution of averaged phase lags between jaw movement rhythmicity and theta oscillations. Top, polar plot of averaged phase lags for individual units, with radial axes representing Rayleigh z scores. Bottom, distributions of phase lags for significantly and non-significantly phase-locked units (Rayleigh test, $p < 0.05$). **(E)** Relationship between jaw-rhythm phase-locking strength and spatial selectivity. Jaw phase-locking strength was quantified using Rayleigh z scores. Each dot represents one unit. Correlation was estimated using linear regression with 95% confidence intervals and compared with null distributions generated from 2,000 spike-shuffled controls. *$p < 0.05$, **$p < 0.01$, ***$p < 0.001$. See Supplementary Table 1 for statistical results.

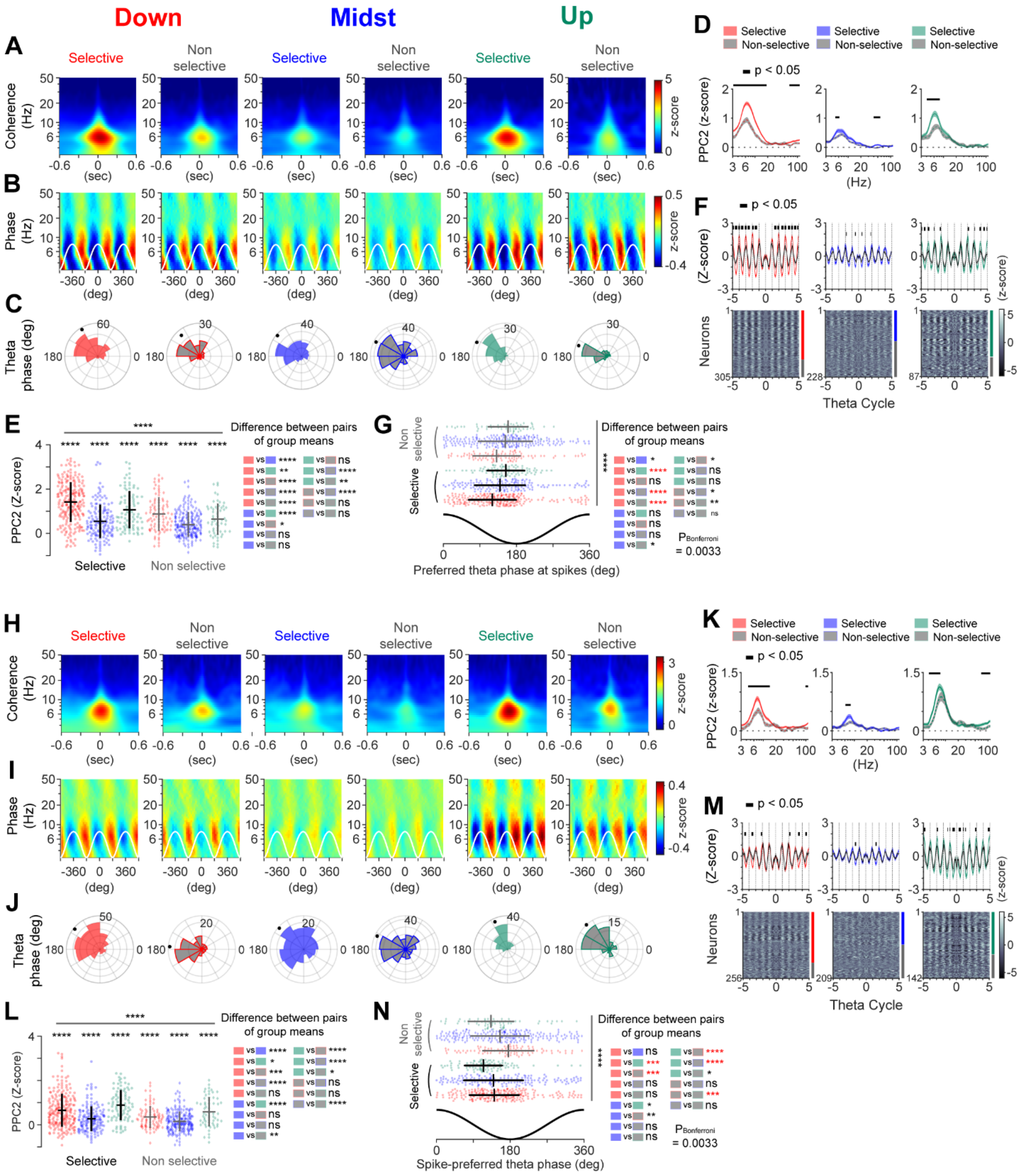

### Supplementary Figure 9. Theta-spike synchronization during the 5 s following chewing onset and the 5 s preceding forepaw contact (related to Figure 5).

**(A)** Population-averaged spike-LFP coherence during the 5 s following chewing onset. For each unit, wavelet-based spike-LFP coherence was z-transformed using 500 shuffled spike trains. Warmer colors indicate stronger spike-LFP coherence at a given frequency and time lag. Spatially selective units exhibited stronger synchronization with ~6-Hz oscillations across reward subtypes.

**(B)** Population-averaged spike-phase distribution maps. Spike-phase histograms were z-transformed using 500 shuffled spike trains. Two theta cycles are shown for visualization. **(C)** Population distributions of preferred theta phases (5–8 Hz). Each dot on the polar axis represents the circularly averaged preferred phase of one population. **(D)** Population-averaged pairwise phase consistency (PPC2) spectrograms showing differences between spatially selective and non-selective units. PPC2 values were z-transformed using 500 shuffled spike trains and are shown as mean ± s.e.m. Horizontal bars indicate significant frequency ranges identified by cluster-based permutation testing (two-sided, $p < 0.05$). **(E)** Beeswarm plots of PPC2 values in the theta range (5–8 Hz) for individual units. Statistical comparisons were performed using paired t-tests within each reward subtype and spatial selectivity category. Mean PPC2 values across subtypes were compared using one-way ANOVA with post hoc multiple-comparison correction. *$p < 0.05$, **$p < 0.01$, ***$p < 1e{-}3$, ****$p < 1e{-}4$. **(F)** Top, population-averaged spike-phase autocorrelograms (mean ± s.e.m.) with significant intervals identified by cluster-based permutation testing (two-sided, $p < 0.05$). Bottom, spike-phase autocorrelogram heatmaps for individual units. Persistent phase locking was prominent in spatially selective ramp-down and ramp-up units. **(G)** Beeswarm plots showing preferred theta phases for individual units. Statistical comparisons were performed using the Watson-Williams test with post hoc multiple-comparison correction. Red asterisks indicate significance exceeding the Bonferroni-corrected threshold. **(H–N)** Same as A–G, but for the 5 s preceding forepaw contact. See Supplementary Table 1 for statistical results.

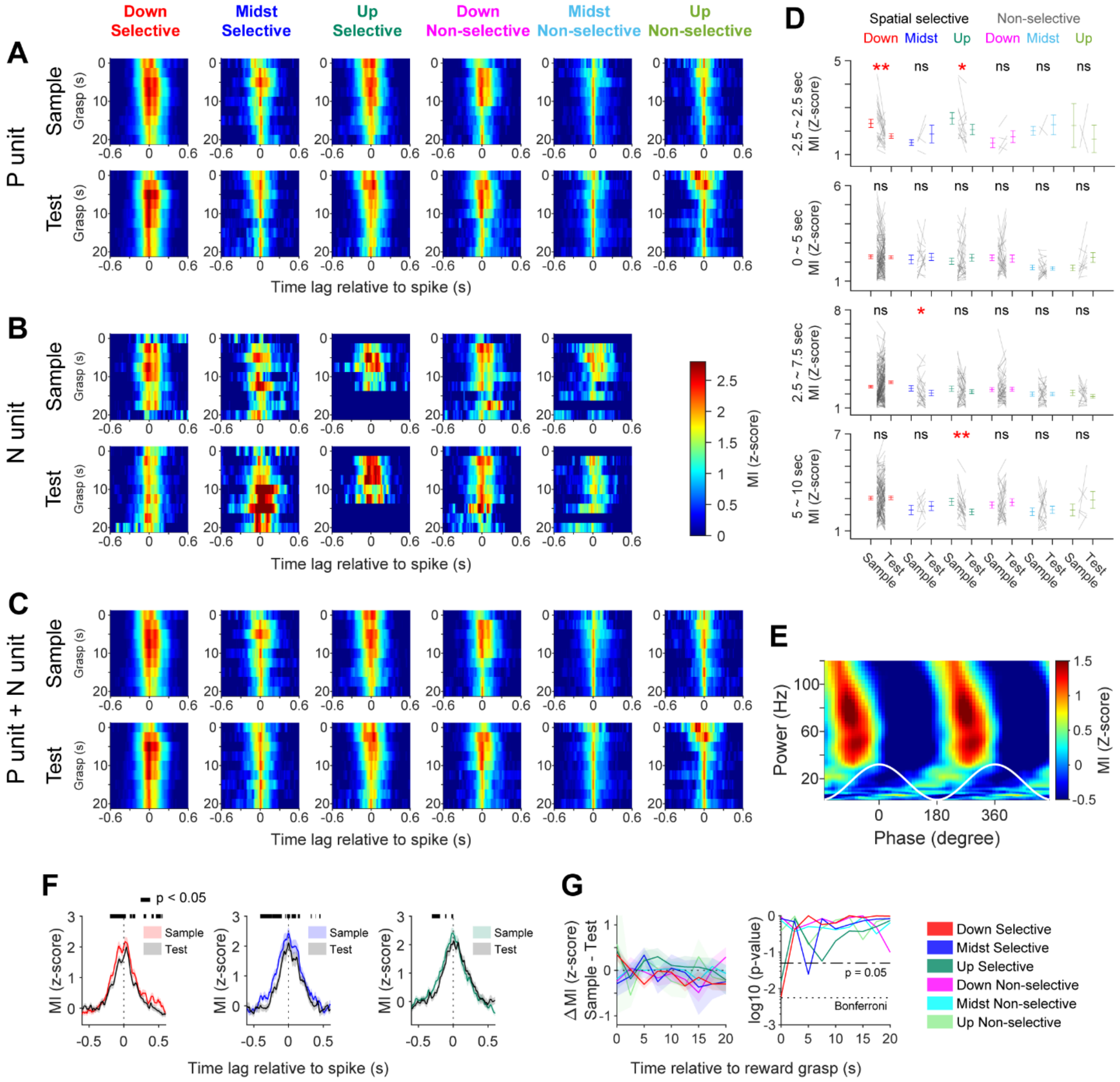

Supplementary Figure 10. Trial-dependent theta-high gamma phase-amplitude coupling at spikes during the initial phase of consummation (related to Figure 6).

**(A–C)** Population-averaged, time-resolved spike-triggered TG-PAC in the high-gamma range (60–120 Hz). Top, sample trials; bottom, test trials. Sample and test trial numbers were equalized. TG-PAC was computed using 5 s bins with 2.5 s sliding windows, and coupling strength was quantified as z-scored spike-triggered coupling relative to spike-shuffled surrogate datasets (100 shuffles). (A) P-units. (B) N-units. (C) P-units and N-units pooled together. **(D)** Comparison of spike-triggered TG-PAC between sample and test trials for individual units within each reward subtype. Paired t-tests were performed within the first four 5 s sliding windows. Black lines represent individual units, and error bars indicate population means ±1 s.e.m. One-sided t-tests were used (*$p < 0.05$, **$p < 0.01$). **(E)** Heatmap of LFP-based TG-PAC during the consummatory period. The x-axis represents theta phase (two cycles shown for visualization), and the y-axis represents gamma-band frequencies. Coupling strength was quantified as z-scored TG-PAC relative to surrogate datasets generated by circularly shifting theta-phase time series relative to gamma-amplitude time series (500 shuffles). Warmer colors indicate stronger theta-gamma coupling. **(F)** Population-averaged peri-spike TG-PAC histograms for spatially selective ramp-down, midst, and ramp-up units comparing sample (colored) and test trials (black). Significant peri-spike intervals were identified using cluster-based paired permutation testing (500 shuffles, one-sided, $p < 0.05$). **(G)** Population-averaged, time-resolved spike-triggered TG-PAC. Top, difference in TG-PAC between sample and test trials (sample − test) for each subtype. Bottom, corresponding log10-transformed p values from statistical comparisons across time. See Supplementary Table 1 for statistical results.

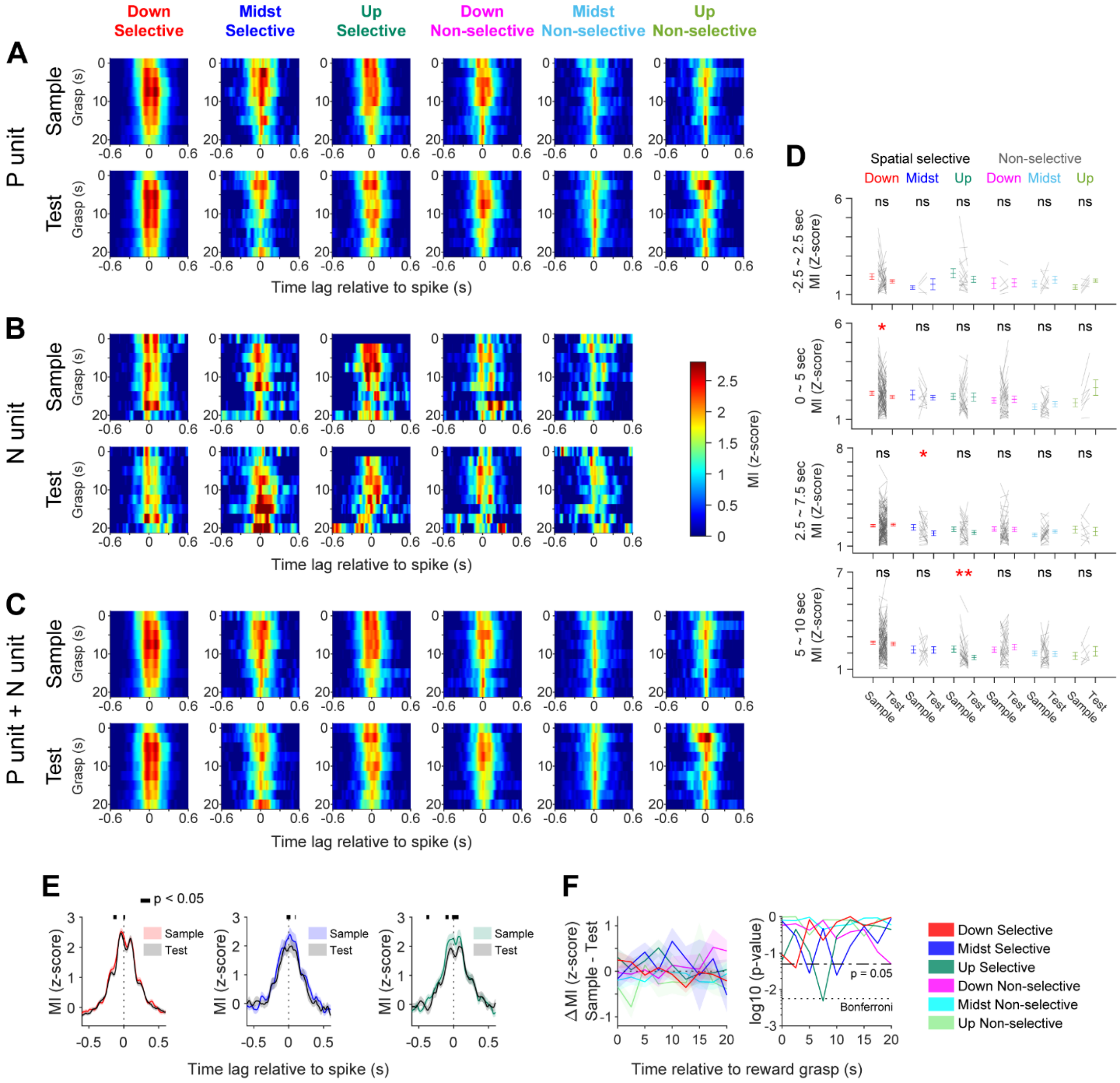

### Supplementary Figure 11. Trial-dependent theta-low gamma phase-amplitude coupling at spikes during the initial phase of consummation (related to Figure 6).

Analysis and plotting procedures were the same as in Supplementary Figure 10, except that TG-PAC was computed in the low-gamma range (30–55 Hz). See Supplementary Table 1 for statistical results.

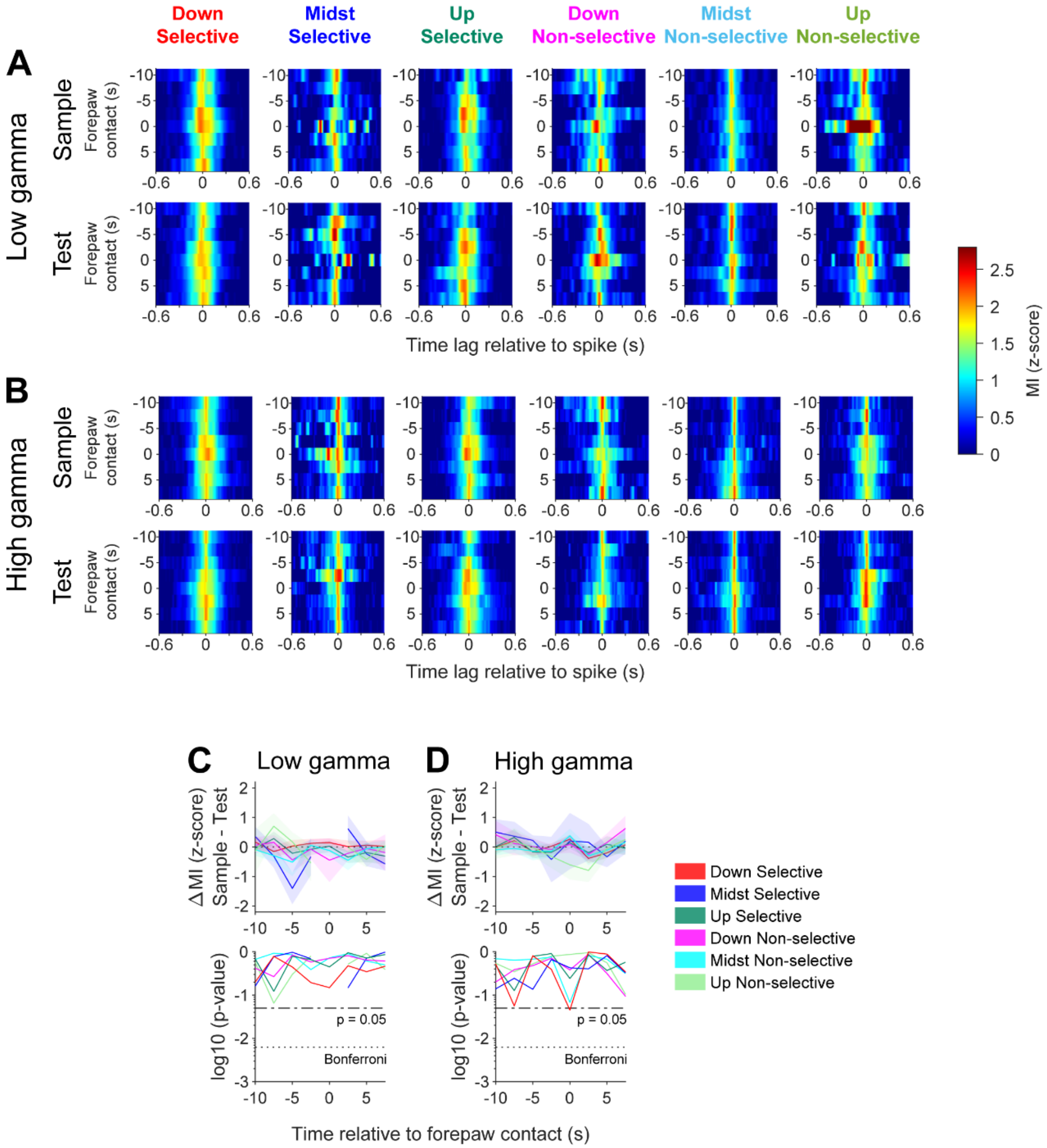

Supplementary Figure 12. Trial-dependent theta-gamma phase-amplitude coupling at spikes during the terminal phase of consummation (related to Figure 6).

**(A,B)** Population-averaged, time-resolved spike-triggered TG-PAC during the terminal phase of consummation, computed using 5 s bins with 2.5 s sliding windows. Top, sample trials; bottom, test trials. Sample and test trial numbers were equalized. Coupling strength was quantified as z-scored spike-triggered coupling relative to spike-shuffled surrogate datasets (100 shuffles). P-units and N-units were pooled together. (A) Low-gamma range. (B) High-gamma range. **(C,D)** Population-averaged, time-resolved spike-triggered TG-PAC. Top, difference in TG-PAC between sample and test trials (sample − test) for each subtype. Bottom, corresponding log10-transformed p values from statistical comparisons across time. (C) Low-gamma range. (D) High-gamma range.

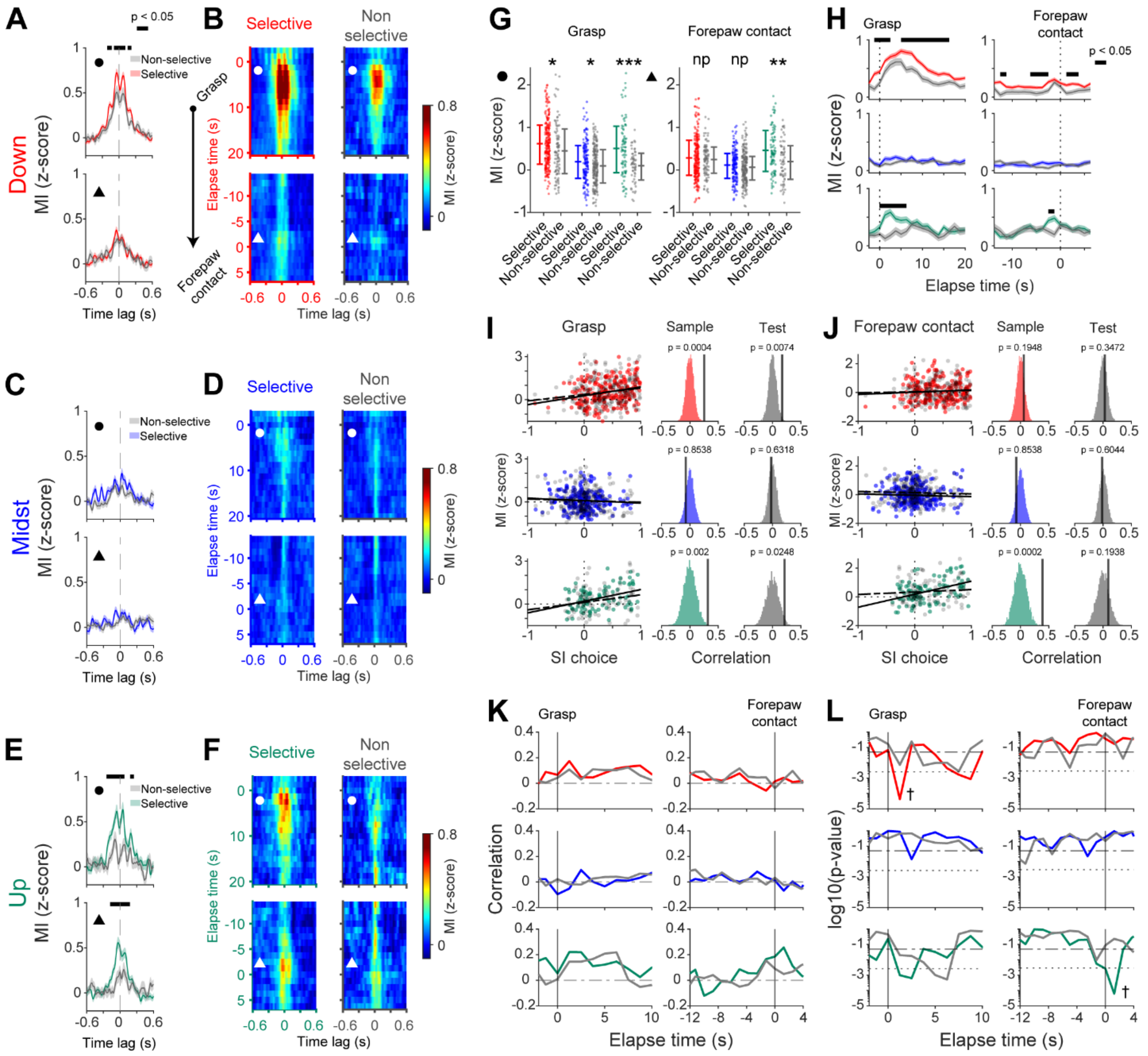

**Supplementary Figure 13. Trial-dependent modulation of low-gamma TG-PAC at reward-unit spikes reflects spatial encoding during DNMP T-maze working memory task (related to Figure 6).** **(A,C,E)** Population-averaged peri-spike TG-PAC in the low-gamma range (30–55 Hz) for spatially selective and non-selective ramp-down (A), midst (C), and ramp-up (E) units. Time 0 indicates spike timing. Black horizontal lines indicate significant differences identified by cluster-based permutation testing (500 shuffles, two-sided, $p < 0.05$). Top, 2.5 s epochs aligned to reward grasp; bottom, 2.5 s epochs aligned to forepaw contact. Reward units were included regardless of negative TG-PAC modulation. **(B,D,F)** Time-resolved population-averaged low-gamma TG-PAC for ramp-down (B), midst (D), and ramp-up (F) units. Circles and triangles indicate the time bins used for peri-spike TG-PAC analyses shown in A, C, and E. **(G)** Top, beeswarm plots of TG-PAC values for spatially selective and non-selective units across reward subtypes during the 2.5 s period following reward grasp. Statistical differences were assessed using unpaired t-tests (ramp-down: $t(326) = 2.44$, $p = 1.51e{-2}$; midst: $t(347) = 2.03$, $p = 4.32e{-2}$; ramp-up: $t(140) = 4.29$, $p = 3.25e{-4}$).

Bottom, same analysis for the 2.5 s period preceding forepaw contact (ramp-down: $t(322) = 6.74e{-1}$, $p = 5.01e{-1}$; midst: $t(359) = 1.32$, $p = 1.87e{-1}$; ramp-up: $t(156) = 3.34$, $p = 1.03e{-3}$). *p <5e−2, **p < 1e−2, ***p < 1e−3. **(H)** Time-resolved population-averaged TG-PAC for spatially selective and non-selective units. Significant intervals were identified separately for each reward subtype using cluster-based permutation testing (500 shuffles, two-sided, $p < 0.05$). **(I)** Top, relationship between spatial selectivity index and spike-triggered TG-PAC during sample and test trials. Each dot represents one unit; lines indicate linear regression fits. Kendall's tau correlations were computed for each subtype (ramp-down sample: $\rho = 0.25$, $p = 6.28e{-5}$; ramp-down test: $\rho = 0.17$, $p = 5.35e{-4}$; midst sample: $\rho = -7.44e{-2}$, $p = 0.87$; midst test: $\rho = -2.43e{-2}$, $p = 0.64$; ramp-up sample: $\rho = 0.32$, $p = 1.15e{-4}$; ramp-up test: $\rho = 0.21$, $p = 2.14e{-2}$). Middle, sample-trial correlations compared with null distributions generated using 5,000 spike shuffles per unit. Bottom, same analysis for test trials. **(J)** Same as I, but for the 2.5 s period aligned to forepaw contact. **(K)** Time-resolved Kendall's tau correlation between spatial selectivity index and peri-spike TG-PAC. Colored lines indicate sample trials; gray lines indicate test trials. **(L)** Time-resolved p values for Kendall's tau correlations. Dashed horizontal lines indicate Bonferroni-corrected significance thresholds; dagger symbols indicate time points exceeding corrected significance levels.

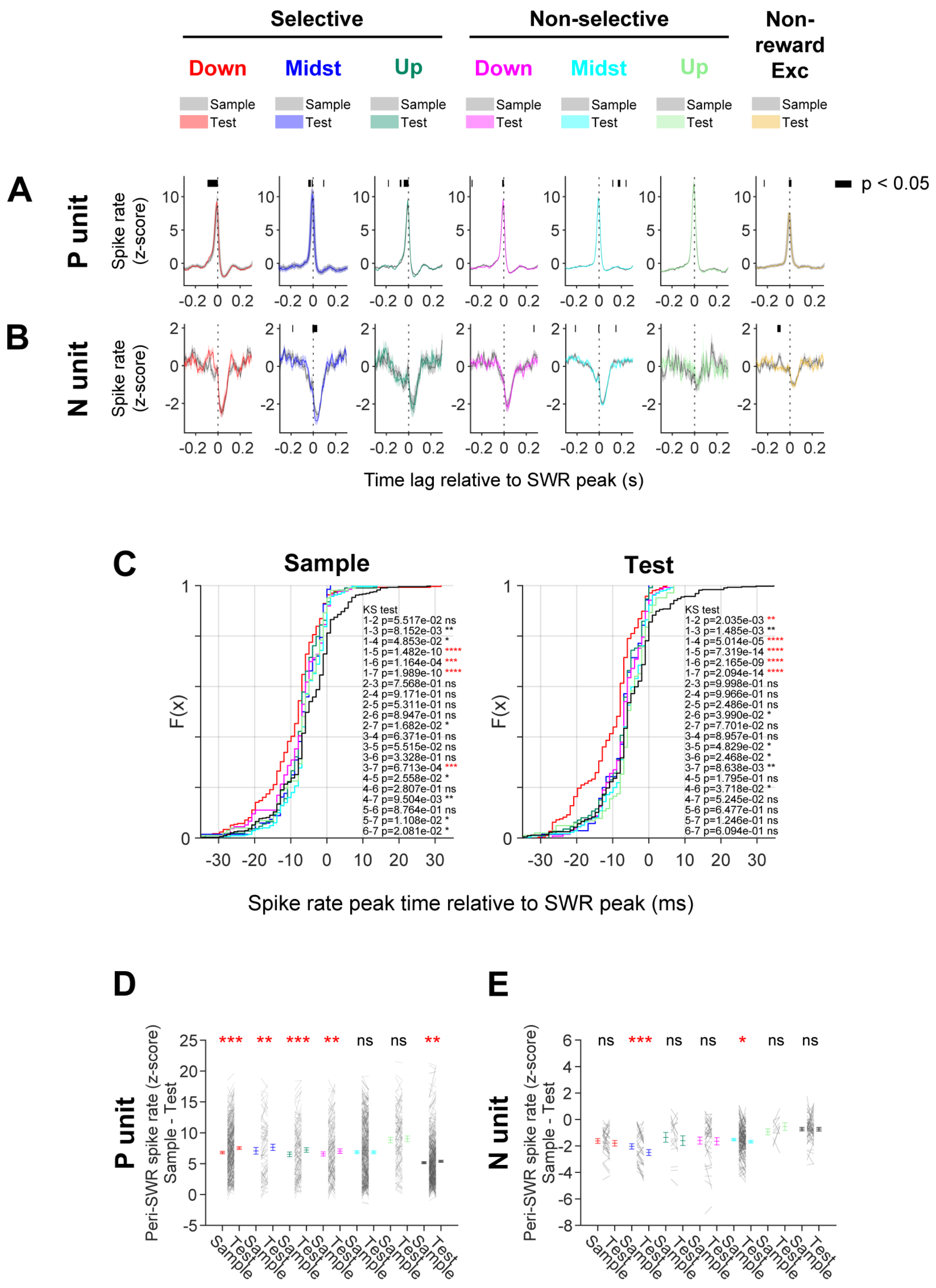

Supplementary Figure 14. Trial-dependent SWR modulation of reward units during consummation at reward sites ipsilateral to each unit's theta field (related to Figure 7).

**(A)** Population-averaged peri-SWR spike rates (z-scored) for P-units. Spatially selective (ramp-down, midst, ramp-up), spatially non-selective (ramp-down, midst, ramp-up), and non-reward excitatory units are shown with shaded areas representing ±1 s.e.m. Sample and test trials were compared at reward sites ipsilateral to each unit's theta-related field. Horizontal black bars indicate significant peri-SWR intervals identified by cluster-based paired permutation testing (sample < test, one-sided, $p < 0.05$). **(B)** Same as A, but for N-units exhibiting suppression during SWRs. Horizontal black bars indicate significant peri-SWR intervals identified by cluster-based paired permutation testing (sample > test, one-sided, $p < 0.05$). **(C)** Cumulative distributions of peri-SWR spike peak times relative to SWR peaks during ipsilateral sample trials (left) and test trials (right). Distributions are shown separately for spatially selective and non-selective reward P-units and non-reward excitatory P-units. Insets show p values from pairwise Kolmogorov-Smirnov (KS) tests. Red asterisks indicate comparisons remaining significant after Bonferroni correction. Numeric labels denote subtypes: 1, spatially selective ramp-down; 2, non-selective ramp-down; 3, spatially selective midst; 4, non-selective midst; 5, spatially selective ramp-up; 6, non-selective ramp-up; 7, non-reward excitatory units. **(D)** Peri-SWR peak spike rates for P-units during sample and test trials at ipsilateral reward sites. Black lines connect individual units across trial types. Colored error bars indicate population means ±1 s.e.m. Statistical comparisons were performed using paired t-tests (sample < test, one-sided, *$p < 0.05$, **$p < 0.01$, ***$p < 0.001$). **(E)** Same as D, but for N-units. Statistical comparisons were performed using paired t-tests (sample > test, one-sided). See Supplementary Table 1 for statistical results.

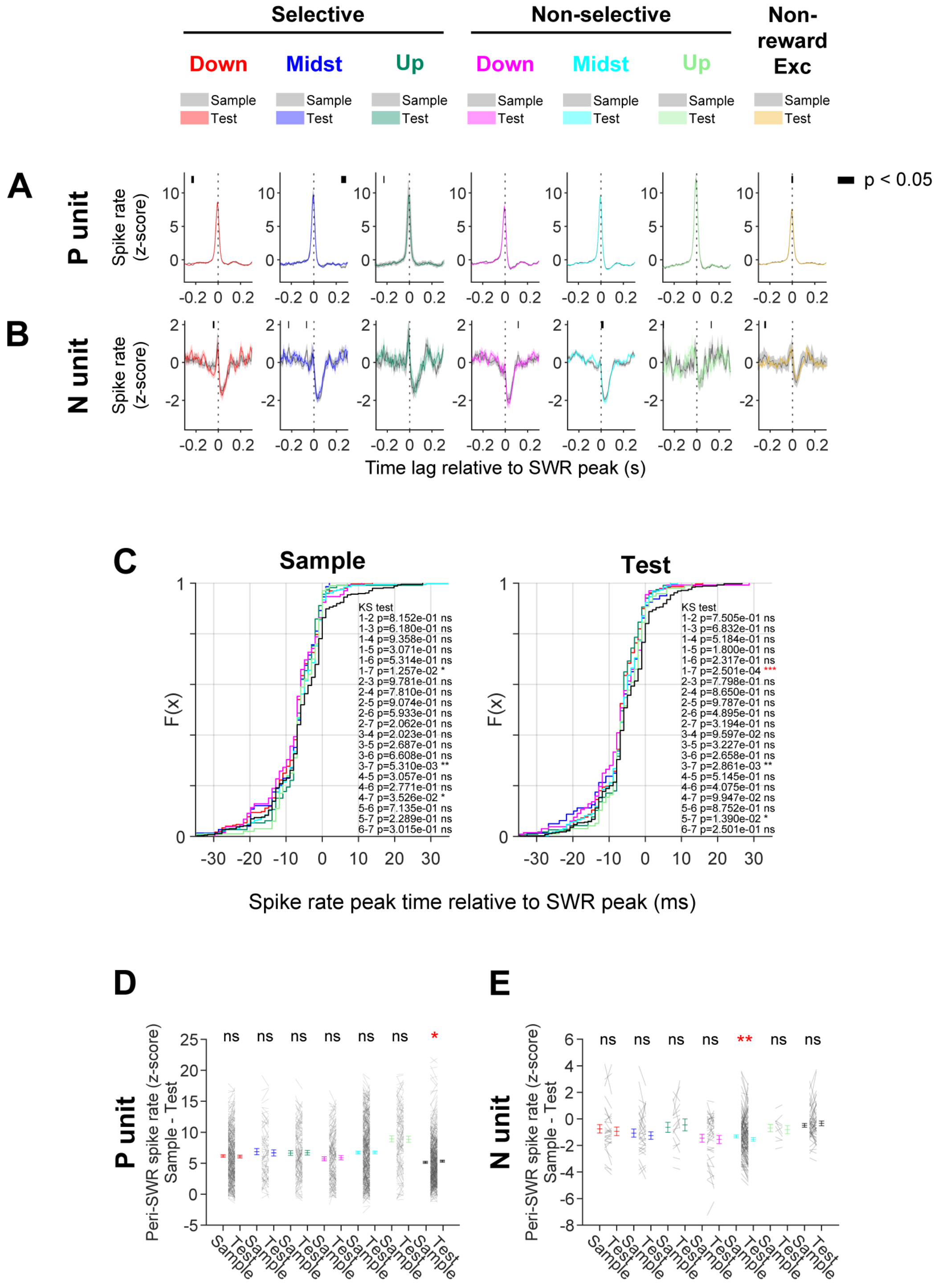

Supplementary Figure 15. Trial-dependent SWR modulation of reward units during consummation at reward sites contralateral to each unit's theta field (related to Figure 8).

**(A)** SWR-triggered spike rates (z-scored) for P-units at contralateral reward sites. Same format as in Supplementary Figure 14A, but for contralateral reward sites. **(B)** SWR-triggered spike rates (z-scored) for N-units at contralateral reward sites. Same format as in Supplementary Figure 14B, but for contralateral reward sites. **(C)** Cumulative distributions of peri-SWR spike peak times at contralateral reward sites. Same format as in Supplementary Figure 14C, but for contralateral sample and test trials. **(D)** Peri-SWR peak spike rates for P-units at contralateral reward sites. Same format as in Supplementary Figure 14D, but for contralateral sites. **(E)** Peri-SWR peak spike rates for N-units at contralateral reward sites. Same format as in Supplementary Figure 14E, but for contralateral sites. See Supplementary Table 1 for statistical results.

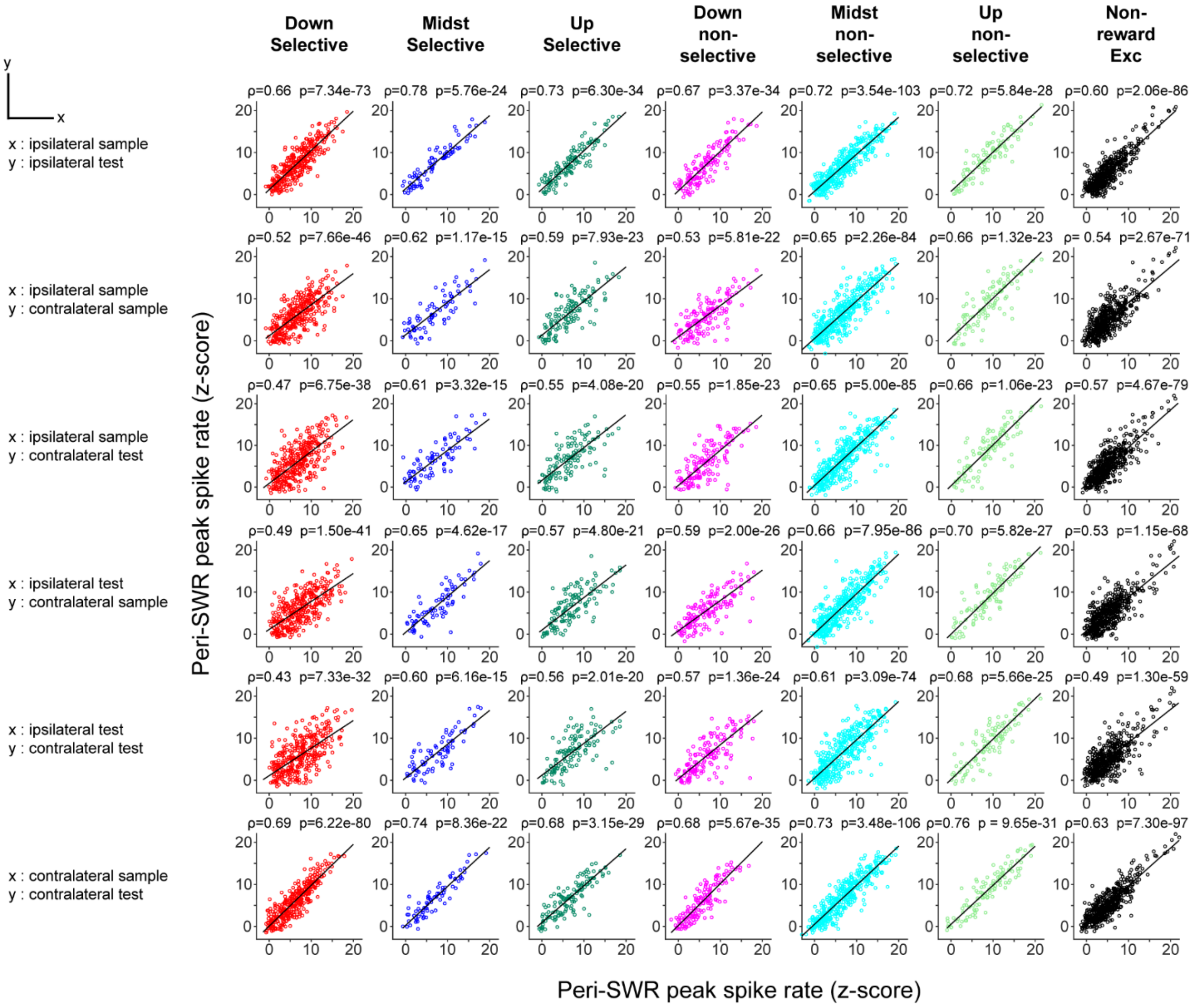

Supplementary Figure 16. Pairwise comparisons of peri-SWR peak spike rates during consummation across trial types and reward sites (related to Figures 7 and 8).

Scatter plots comparing peri-SWR peak spike rates (z-scored) during consummation across trial types and reward sites for individual units. Each dot represents one unit. Columns correspond to spatially selective (ramp-down, midst, ramp-up), spatially non-selective (ramp-down, midst, ramp-up), and non-reward excitatory units. Rows show all pairwise comparisons among ipsilateral sample, ipsilateral test, contralateral sample, and contralateral test conditions. Pearson correlation coefficients (r) and corresponding p values are shown for each comparison.

### Supplementary Tables

Supplementary Table 1 (Provided as a separate Excel file)
